# Sex difference in the underpinnings of social fate in isogenic mice and no sex bias in precedence-based social ranking in male and female groups

**DOI:** 10.1101/2025.06.25.660781

**Authors:** Estelle Conabady, Cécile Vernochet, Aylin Gulmez, Fabio Marti, Philippe Faure, Sébastien Parnaudeau, François Tronche

## Abstract

The mechanisms underlying social organization in mice have predominantly been studied in male colonies, where sociability predicts higher rank and high-ranked individuals show greater anxiety. Isogenic female mice also form stable social hierarchies. Our data indicate that greater sociability before group formation predicts high rank attainment, a trait that remains afterwards. In contrast, a consistent increased anxiety was only observed after life in colony for the same individuals. Mixed-sex colonies, revealed a similar hierarchical structure, with both males and females having equal chances of attaining high and low ranks. We previously found reduced dopamine neuron activity in highest-ranked males. In contrast, highest-ranked females show the opposite pattern. Furthermore, whereas in males, glucocorticoid receptor signaling impairment in dopaminoceptive neurons favored high rank attainment, this effect is absent in females. Overall, these results highlight sex-specific mechanisms that contribute to social ranking and related behavioral traits in mice.

## Introduction

The emergence of social organization is a widespread phenomenon in the animal kingdom, offering several evolutionary advantages, including the reduction of inter-individual conflicts, task allocation, cooperation and more efficient resource sharing ^1,2^. At the individual level, social status can contribute to the development of individual behavioral traits that reflect diverse strategies for foraging, predator avoidance, and behavioral adjustments with conspecifics. Social hierarchy is a flexible structure influenced by factors such as colony size, food availability, and territorial boundaries. In the wild, house mice (*Mus musculus*) inhabiting large natural environments with limited resources tend to be less territorial and adopt a sedentary lifestyle only during reproduction and pup-rearing periods ^3^. In contrast, commensal mice living alongside humans form smaller social groups of four to eight adults, usually with one dominant male and subordinate individuals that may be either males or females ^4,5,6^.

Over the past decade, laboratory research has intensified effort to understand the physiological mechanisms and brain circuits underlying the emergence of social organization and its impact on behavioral individuation. Social organization of rodents has been studied in standard laboratory cages ^7, 8^, semi-naturalistic environment ^9, 10, 11^, and large buildings or a gardens ^12, 13^. In these different contexts, social ranking can be assessed through observations of agonistic interactions, including threats, chases, and attacks ^7–15^; territorial urine-marking ^7, 16^, and territorial occupancy ^7–14^; access to limited resources, such as food and water ^7, 8, 17–19^ or occupancy of a warm spot in a cold cage ^20, 21^; and precedence behaviors measured in the _Tube-test 7, 8, 16, 18-24._

The emergence of social organization in rodents has been studied almost exclusively in same-sex colonies, predominantly in males. One reason may be the difficulty assessing dominance-related behaviors commonly used to evaluate hierarchy in males, such as urine marking, vocalization, and overt aggression. A growing body of evidence indicates that female mice establish stable social hierarchies, although these are generally less despotic and less linear than those observed in males ^11, 23–33^. This difference may reflect behavioral variability associated with the estrus cycle, together with the marked reduction of aggressivity in laboratory mouse females compared with wild mouse females. In semi-naturalistic environments, groups of wild female mice form hierarchies that are as robust and structured as those of males ^11^. Marked sex differences in social organization and competitive strategies have been reported between male and female colonies ^11, 26^, as have differences in the underlying physiological mechanisms, including sexually dimorphic circuits ^19, 32^ and long-term potentiation at synapses connecting the mediodorsal thalamus to the dorsomedial prefrontal cortex ^33^. Only a few studies have examined intersexual social ranking in laboratory rodents, and in mixed-sex colonies, social rank has often been assessed separately within each sex. ^9,10.^

We previously analyzed social ranking in colonies of four isogenic C57Bl/6 male mice using the precedence Tube-test. We previously demonstrated that highest ranked individuals exhibit enhanced anxiety and sociability compared with their lower-ranked counterparts. While differences in anxiety emerged after colony formation, increased sociability in future high-ranking individuals was already evident prior to the establishment of social hierarchy. At the physiological level, we found that reducing dopamine neuron activity in the ventral tegmental area (VTA) facilitates access to higher social rank, an effect that was mimicked by genetically inactivating the glucocorticoid receptor (GR) in dopamine-receiving neurons ^21^.

To address the determinants of social hierarchy formation in female laboratory mice, we adopted the same experimental design as in our previous work on male mice. We observed that C57BL/6 female mice housed in groups of four, rapidly develop a stable social hierarchy. By assessing individual behavior before and after colony formation, we found that increased sociability was a pre-existing trait in females that ultimately attain the highest rank. In contrast, a consistent increased anxiety-like behavior in the same individuals was only observed after life in colony for the same individuals. To examine whether hierarchical ranking is influenced by sex, we formed mixed-sex colonies by housing a vasectomized male with three females. These groups established a hierarchical structure similar to that of unisex colonies, with both sexes showing an equal likelihood of attaining either the highest or lowest rank. Females exhibited an opposite pattern of VTA dopamine neurons activity, increased among high-ranking individuals, and social rank attainment was not affected by glucocorticoid receptor signaling in dopaminoceptive neurons.

## Methods

### Animals

C57BL/6JRj, female mice, aged six, eight, and nine weeks and 6-week-old vasectomized male mice were purchased from Janvier Labs (Le Genest-Saint-Isle, France). Mice were maintained under standard laboratory conditions (22°C, 55% to 65% humidity) on a 12h light/dark cycle (lights on at 07:00) and had *ad libitum* access to food and water. All procedures complied with the European Directive 2010/63/EU and Recommendation 2007/526/EC for animal care and use, and were approved by the Sorbonne Université animal ethics committee.

Selective inactivation of the *Nr3c1* (GR) gene in dopaminoceptive neurons was achieved using *Nr3c1*^loxP*/*loxP^;Tg:D1aCre mice (hereafter referred to as GR^D1aCre^), as previously described by Ambroggi *et al.* 2009 ^34^. Experimental animals were generated by crossing Nr3c1^loxP*/*loxP^ females with Nr3c1^loxP*/*loxP^;Tg:D1aCre males, yielding approximately equal proportion of mutant and littermate controls.

For C57BL/6 female tetrads, upon arrival, mice were weighed and assigned to groups of four weight-matched individuals, hereafter referred to as tetrad. If a first behavioral testing preceded group formation, mice were singly housed for seven to ten days, until grouping during which the tests were performed (see Fig. S1). In mixed-sex tetrads, one vasectomized male was housed with three females, aged around nine and eighteen weeks old, respectively. Starting at seven and fifteen weeks, respectively, male and female were weighed daily in order to group weight-matched individuals.

To assess the role of GR in dopaminoceptive neurons in social rank, tetrads consisted of one GR^D1aCre^ mutant female and three age- and weight-matched control females (GR^loxP/loxP^), from different litters, unfamiliar to one another. GR^Forebrain^

### Behavioral studies

The timing of the behavioral studies performed in the different tetrads is depicted in Figure S1. For all behavioral studies (see below) and prior to each experiment, mice were accustomed to the experimental room for one hour prior to each experiment. Apparatuses were cleaned with 20% ethanol between subjects to eliminate olfactory cues. For ranked animals, behavioral studies began the week after rank determination and lasted seven to nine days. Data were analyzed taking into account this ranking assignment. All behavioral experiments were scored by an experimenter blind to the experimental conditions. Except for the Tube-test, all sessions were video-recorded and analyzed offline.

### Social rank assessment, Tube test

Following two weeks of group housing, mice were trained to walk through a transparent Plexiglas tube (diameter, 2.5 cm; length, 30 cm) for two consecutive days (8 trials on day 1, 4 trials on day 2). Each mouse entered from alternating ends and was allowed 30 seconds to traverse the tube; mice that failed to do so were gently guided out. The tube diameter allowed only one mouse to pass and prevented turning. Social rank was determined through all six pairwise combinations within each tetrad across three daily repeated sets of trials, each set consisting of three confrontations, *i.e.* nine confrontations per day between two individuals. Two mice were introduced simultaneously from opposite ends of the tube. The mouse that backed out first was considered the loser. Ranks were assigned from 1 to 4 according to the number of confrontations won. Trials lasting longer than 2.5 minutes were aborted and repeated. The order of pairings followed a randomized round-robin schedule.

Ranks were initially assessed over a minimum of six days and considered stable when ranks 1 and 4 remained unchanged for three consecutive days. Tetrads not meeting this criterion were tested further, up to a maximum of 14 days, at which point all repeated Tube-test measurements had reached stability. Additional behavioral tests were performed immediately after rank determination within each tetrad. One cohort underwent repeated testing every month for approximatively four months.

### Warm spot occupancy assay

Tetrads were placed in a transparent plastic cage (35×20×18 cm) without litter and placed on ice (bottom cage temperature 4 °C). 20 minutes later, a warm plate (4.5 x 5 cm, 28-30 °C) was introduced on the floor of the cage, at a corner. Mice activity was recorded for 20 min, and warm plate occupancy by each individual was scored by blocks of 5 min.

### Anxiety-like behavior

#### Dark-light box

Mice were placed in a two-chamber apparatus (45 × 20 × 25 cm) consisting of a lit compartment (30 cm, 200–280 lux) and a dark compartment (15 cm, covered), connected by a 5 × 5 cm opening. Mice were initially placed in the dark compartment, and exploration was recorded for 10 minutes.

#### Elevated O-maze

A circular black PVC maze (5.5 cm width, 56 cm outer diameter) was elevated 30 cm above the floor and divided into four sections—two open and two enclosed by 15.5 cm walls. Mice were placed in a closed section the head directed inward under 200–300 lux, and behavior was recorded for 10 minutes. A mouse was considered in a section when all four paws were within it.

### Despair, forced swim test

Mice were placed in a transparent cylinder (40 cm height, 12 cm diameter) filled with water (23°C, 10 cm depth) and recorded for 6 minutes. Behavior was scored as follows: (i) escape behavior, consisting of vigorous wall-climbing movements involving all four paws; (ii) Balance movements: minimal locomotion using primarily the hind limbs; iii) Immobility: floating with no purposeful movement. The test was repeated 24 hours later if required by the experimental design.

### Sociability, three-chambers test

The apparatus consisted of a rectangular box divided into three chambers (30 × 20 × 15 cm) connected by 5 × 5 cm doorways at the center of each partition, under 50-30 lux illumination. Transparent perforated containers (10 × 7 × 7 cm) were placed in the two side chambers, one containing an unfamiliar adult male C57BL/6 mouse and the other left empty. Mice were habituated for five minutes in the central chamber with the doors closed. The doors were then opened, and mice were allowed to explore freely for five minutes. Interaction time, defined as head-directed, close proximity contact, was quantified.

To assess social memory, the test was extended as follows: the mouse was first confined with the social stimulus for five minutes and was then allowed to explore the apparatus again. Time spent interacting with the familiar versus a novel unfamiliar mouse was scored during a second five minute session.

### Locomotor activity

Mice were placed for two hours in a circular transparent PVC corridor (4.5 cm wide, 17 cm external diameter) equipped with four infrared sensors positioned 1.5 cm above the base at quarter intervals (Cyclotron, iMetronics, Bordeaux). Locomotor activity was recorded automatically.

### *In vivo* electrophysiological recordings

Highest and lowest ranked female mice, hereafter referred to as rank 1 (R1) and rank 4 (R4), aged 3 to 5 months, were anesthetized with chloral hydrate (8%, 400 mg kg^−1^ *i.p.*) and placed in a stereotaxic frame. A hole was drilled in the skull above midbrain dopaminergic nuclei (coordinates: 3.0 ± 1.5 mm posterior to bregma, 1 ± 1 mm lateral to the midline). Recording electrodes were pulled from borosilicate glass capillaries using a Narishige electrode puller. Electrode tips were broken under microscopic control and filled with 0.5% sodium acetate. Electrodes had tip diameters of 1-2 µm and impedances of 20–50 MΩ. A reference electrode was placed in the subcutaneous tissue.

Recording electrodes were lowered vertically through the skull opening with a microdrive. Electrical signals were amplified using a high-impedance amplifier and monitored with an oscilloscope and an audio monitor. Unit activity was digitized at 25 kHz and stored using Spike2 software. Electrophysiological characteristics of dopamine neurons were analyzed from active cells encountered during systematic electrode passes through a stereotaxically defined brain region encompassing the VTA. The sampled region extended from -2.9 to -3.5 mm posterior to bregma (AP), 0.3 to 0.6 mm lateral (ML) and -3.9 to -5 mm ventral (DV) ^35^. Recordings were performed first on the right side and then on the left side. Extracellular identification of dopamine neurons was based on their anatomical location and on the set of unique electrophysiological properties distinguishing them from non-dopamine neurons *in vivo*: (i) a characteristic triphasic action potential with a pronounced negative deflection; (ii) a long action potential duration (>2.0 msec); (iii) an action potential width from onset to negative trough >1.1 msec; (iv) a slow firing rate (> 1 Hz and < 10 Hz).

Electrophysiological recordings were analyzed using the R software. Dopamine neuron activity was quantified by the mean firing rate and the percentage of spikes within bursts (%SWB), defined as the number of spikes occurring within bursts divided by the total number of spikes. Bursts were identified as discrete events consisting of a sequence of spikes with an onset marked by two consecutive spikes separated by an interval < 80 ms and terminating with an interspike interval > 160 ms. Firing rate and %SWB were calculated using successive of 60 second windows with a 45 second overlap. The Firing rate and %SWB of each recorded neuron are used as independent variable. Normality of data set is test by Shapiro test. For firing frequency, comparison between group is made using Wilcoxon signed-rank test. For %SWB, because the data set exhibit skewed distribution, comparison between group is made using surrogate method. We first calculate the absolute difference of mean %SWB between the two groups (Δ<%SWB>_o_). We then generated 10 000 surrogate data (Δ<%SWB>_S_). A surrogate data is obtained by resampling two groups from the original dataset (by permutation) and for each resampling by calculating the difference in the mean %SWB of the two resampled group. Original dataset is obtained by pooling the %SWB observed in R1 and R4 mice and resampling by randomly re-assign these values to two groups. We then calculate 10 000 surrogate values Δ<%SWB>_S_ and count the number of times Δ<%SWB>_S_ ≥ (Δ<%SWB>_o_. The null hypothesis is that all samples (From R1 and R4) come from the same distribution. The difference reaches statistical significance if the surrogate reproduces the absolute difference of mean %SWB less than 500/100/10 times over 10 000 simulations.

### Statistical Analysis

Data are expressed as mean ± SEM. Statistical analyses included the Mann-Whitney non-parametric test; unpaired t-test; Gehan-Breslow-Wilcoxon test; Wilcoxon matched-pairs signed rank test; unpaired Kruskal-Wallis test; followed by a Dunn’s multiple comparisons test; and two-way repeated measures ANOVA followed by a Bonferroni’s multiple comparisons test. Statistical significance was set at P ≤0.05. Analyses were performed using Prism software (GraphPad). For each panel, statistical parameters are detailed in the joined dataset.

## Results

### Social hierarchy within a colony (tetrad) of female mice

To determine whether female mice, like males, exhibit social hierarchies, we formed colonies (tetrads) of four C57BL/6 females aged 6-9 weeks (Fig. 1 A, Fig. S1). Rat body mass before the formation of social groups influences Tube-test ranking ^18^. Although this has not been observed in mice ^22, 27, 29, 36^, we grouped weight-matched individuals to avoid any interaction. Two weeks later, we assessed individual social rank using the Tube-test based on pairwise encounters within a plastic tube. The lowest-ranked individual is identified by retreating backward out of the tube ^22^ (Fig. 1A). In males, the Tube-test reveals stable social organization and has been reported to correlate with other status-related behaviors, including occupation of a warm spot in a cold environment, patterns of urine marking and outcome in food competition ^7, 8, 16, 18–21, 23^. All six possible pairwise combinations within each tetrad were tested nine times a day. Individuals were ranked according to the number of forward wins, with the mouse showing the highest number classified as the top-ranked animal. We tested each tetrad daily for at least six consecutive days. Testing continued until both the highest rank (rank 1; hereafter R1) and the lowest rank (rank 4; hereafter R4) were occupied by the same individuals for three consecutive days. Across the 33 tetrads analyzed, rank measurement stability was reach significantly faster for R1 an R4 mice than for intermediate-ranked animals (R2 and R3) (R1 vs R2 and R3 P < 0.0001, R4 vs R2 and R3 P < 0.001, Gehan-Breslow-Wilcoxon test Fig. 1B). As previously reported in male mice ^16, 21^, social rank also influenced the contest duration in females. Encounters between R1 and R2 individuals lasted on average 9 seconds whereas confrontations involving R4 individuals were approximately threefold shorter (P < 0.0001 Kruskal-Wallis test, Fig. 1C). We next assessed the long-term stability of social organization within female colonies. The upper panel of Fig. 1D illustrates the social trajectories of individuals from 12 tetrads monitored across four sessions over a 4-month period. Among the 48 individuals, only 13 maintained the same rank throughout the entire experiment (pink circles, Fig. 1D, lower panel). Compared with male colonies described in our previous study, the female hierarchy appeared significantly less stable (Fig. 1D, lower panel). Between sessions 1 and 2, 10 of the 12 individuals initially identified as R1, and 9 of the 12 initially identified as R4 retained their respective ranks (compared with 12 and 11, respectively, in male colonies ^21^). By session 3, these numbers had decreased to 9 R1 and 6 R4 individuals (*vs*. 12 and 11 in males), and by the final session, only 4 R1 and 5 R4 individuals still occupied their original ranks (*vs*. 12 and 10 in males). Notably, at the end of the 4-month period, three individuals initially classified as R1 had shifted to R4, whereas one initially classified as R4 had become R1, further highlighting the pronounced instability of female social hierarchies. Consistent with this observation, comparison of the temporal evolution of rank maintenance between females and males (Fig. 1D, lower panel, pink circles and blue squares, respectively), revealed significant differences for individuals initially ranked as R1 and R4 (Gehan-Breslow-Wilcoxon test, P < 0.05 and P < 0.01, respectively). Since in both cases, we established weight-matched tetrads, differences in weight are unlikely to explain the differences in rank stability between males and females. Of note, we observed that, in contrast to males ^21^, females initially ranked as R1 gained more weight over the four-month period than R4 individuals (Fig. S2). Data were analyzed using repeated two-way ANOVA followed by Bonferroni’s post hoc test. This analysis revealed a significant interaction effect (P < 0.001; F (34, 697) = 2,026) and a strong effect of time (P < 0.0001; F (17, 697) = 359.4), but no overall effect of social rank (P = 0.12; F (2, 41) = 2.251). Post hoc analysis showed that R1 mice were significantly heavier than R4 ones (weeks 14, 19, 21, 23 and 24), and R2/R3 ones (weeks 21 and 24) (Fig.S2).

**Figure 1:**
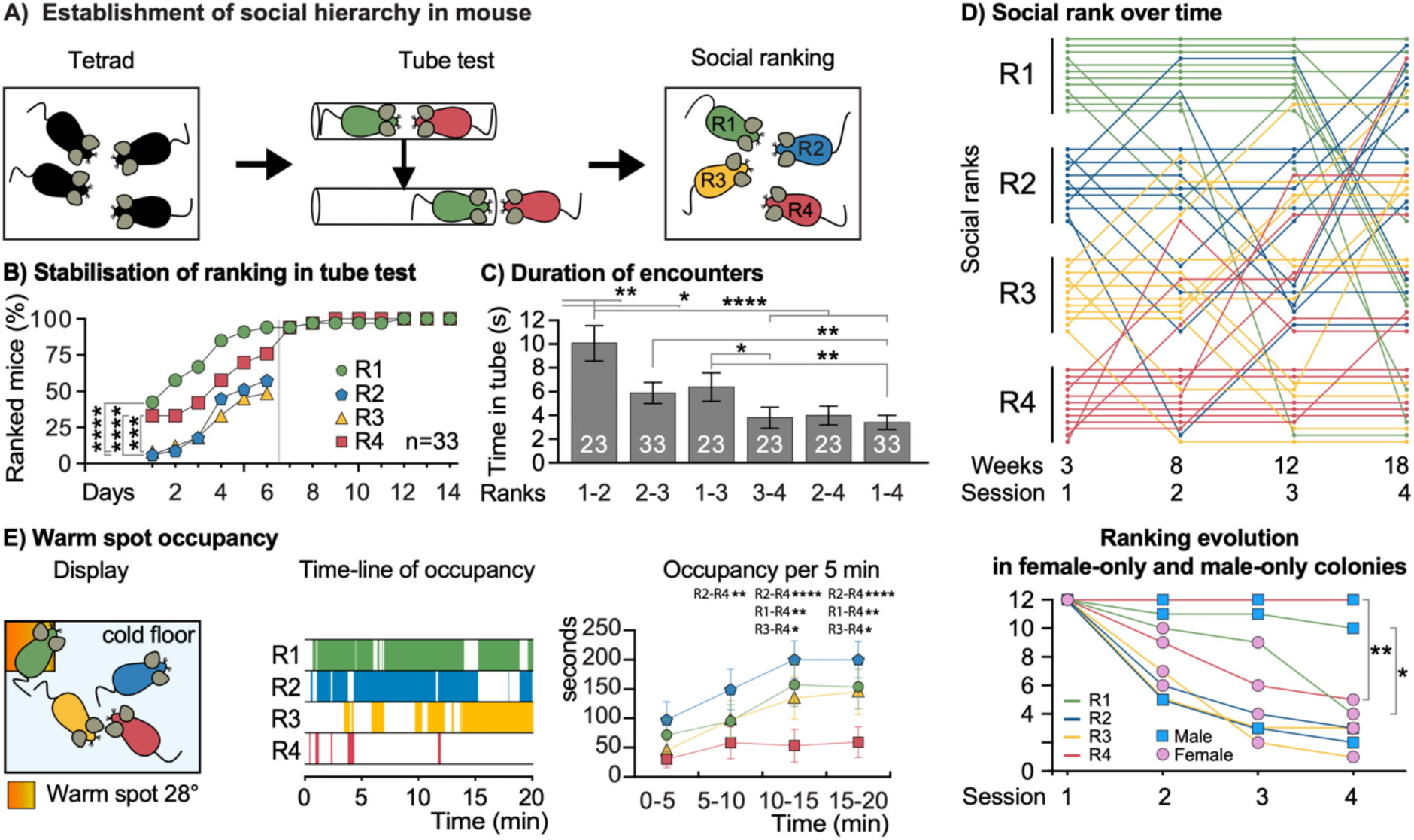
Establishment and stability of social hierarchy in female mice. **(A)** Schematic representation of social hierarchy formation and assessment. Unfamiliar C57BL/6 female mice were grouped into tetrads (groups of four individuals). Two weeks later, their social rank was determined using the Tube-test. **(B)** Dynamics of rank identification in the Tube-test. The cumulative percentage of individuals that reached their final rank is plotted for each testing day (n = 33 tetrads). Results for R2 and R3 individuals are shown only from days 1 to 6 because rank assessment was discontinued after day 6, once R1 and R4 positions had stabilized for three consecutive days. Gehan-Breslow-Wilcoxon test, **** P < 0.0001, *** P < 0.001. Error bars represent ± SEM. **(C)** Average duration of confrontations according to the rank of the opposing individuals, measured during the last three days of social organization stabilization (n = 33 tetrads). All possible rank pairings are represented. Kruskal-Wallis test with post hoc comparison: * P < 0.05; ** P < 0.01; **** P < 0.0001). **(D)** Top panel: Longitudinal evolution of social ranks over 15 weeks in 12 tetrads. Ranks were assessed 3, 8, 12 and 18 weeks after tetrad formation. Each row represents one mouse. Colors indicate the initial rank determined after three weeks: green (R1), blue (R2), orange (R3), and red (R4). Within each rank category, dot position indicates the tetrad to which the individual belonged, from Tetrad 1 (top) to Tetrad 12 (bottom). Changes in rank over time are illustrated by connecting lines. Bottom panel: proportion of mice maintaining stable ranks in female tetrads (pink circles) and male tetrads (blue squares, data from Battivelli *et al.* ^13^). Line colors indicate rank identity. Gehan-Breslow-Wilcoxon test. * P < 0.05, ** P < 0.01. **(E)** Left: schematic representation of the warm spot test, with the heated area indicated in orange Middle panel: representative occupancy periods of the warm spot by individuals from a single tetrad. Right panel: time-course analysis of warm-spot occupancy. Statistical analysis was performed using two-way repeated measures ANOVA followed by Bonferroni’s post hoc test. **** P < 0.0001, ** P < 0.01, * P < 0.05. Error bars represent ± SEM.

To validate the precedence Tube-test as a reliable proxy for social ranking in female mice, we next assessed access to shared but limited resources. Specifically, we quantified occupancy of a warm spot placed within a cold cage positioned on ice by the different individuals of a tetrad during a 20-min period (Fig 1E, left panel) ^20, 21, 33^. Owing to its restricted size (4.5 x 5 cm), the warm spot could hardly accommodate more than two mice simultaneously, as illustrated by the occupancy timeline (Fig 1E middle panel). Using a cohort of 12 tetrads, this assay was performed one day after two consecutive Tube-test sessions (Fig. S3), at both three and nine weeks, following tetrad formation. Data were analyzed using repeated two-way ANOVA followed by Bonferroni’s post hoc test. This analysis revealed a significant effect of time (P < 0.0001; F(3,33) = 23.47) and a significant effect of rank (P < 0.05; F(3,33) = 3.016), with no significant interaction between the two factors (P = 0.5010; F(9,99) = 0.9321). R4 mice identified by the Tube-test spent significantly less time on the warm spot than their cage mates, particularly during the last 10 min of the assay timeline (R4 *vs.* R1, P < 0.01; R4 *vs.* R2, P < 0.001; and R4 *vs.* R3, P <0.05). These findings further support the validity of the Tube-test and confirm the lower social rank of R4 individuals within the hierarchy.

### Social rank correlates with behavioral differences

We next analyzed the relationship between social rank and behavioral phenotypes by comparing R1 and R4 individuals in assays assessing social, anxiety-like, and depression-like behaviors. Behavioral testing was initiated just after Tube-test ranking and was conducted over a period of 7 to 9 days. All analyses were performed according to the established rank assignment.

Sociability was evaluated using the three-chamber test, in which mice choose between exploring a compartment containing an unfamiliar conspecific enclosed in a transparent box and a compartment containing a similar empty box (Fig. 2A, top left). As expected, isogenic C57BL/6 mice (pooled R1 and R4 mice) displayed a clear preference for the social stimulus, though with substantial interindividual variability (Fig. 2A social preference, left middle panel gray bars). Overall, C57BL/6 mice (n = 42) spent significantly more time interacting with the box containing a conspecific than with the empty box (1.07 ± 0.03 *vs.* 1.30 ± 0.06 AU: arbitrary units corresponding to normalized interaction time; respectively; P < 0.001 Wilcoxon matched pairs signed-rank test). Interindividual variability was broad, ranging from 0.69 to 1.49 AU for interaction with the empty box and from 0.07 to 2.08 AU for interaction with the unfamiliar conspecific. Stratification by social rank revealed that R1 individuals displayed a significant social preference (Fig. 2A social preference, left middle panel green bars), whereas R4 mice did not (red bars). Specifically, R1 individuals (n=21) spent 1.00 ± 0.03 AU interacting with the empty box *vs.* 1.29 ± 0.06 AU interacting with the box containing an unfamiliar conspecific. In contrast, R4 individuals (n=21) spent 1.14 ± 0.04 AU *vs.* 1.31 ± 0.07 AU, interacting with the empty and social boxes, respectively. Data were analyzed using two-way repeated-measures ANOVA followed by Bonferroni’s post hoc test. No interaction between social rank and stimulus type was detected (P = 0.38; F(1,40) = 0.78), and no overall effect of social rank was observed (P = 0.1945; F(1,40) = 1.741). However, there was a significant effect of social cue (P < 0.01; F(1,40) = 11.75). Post hoc analysis showed that R1 mice spent significantly more time interacting with the social stimulus than with the empty box (P < 0.01), whereas this difference did not reach significance in R4 mice (P = 0.16). These findings indicate that part of the interindividual variability in sociability observed among female mice can be explained by social rank, as previously described in males ^21, 23^.

**Figure 2:**
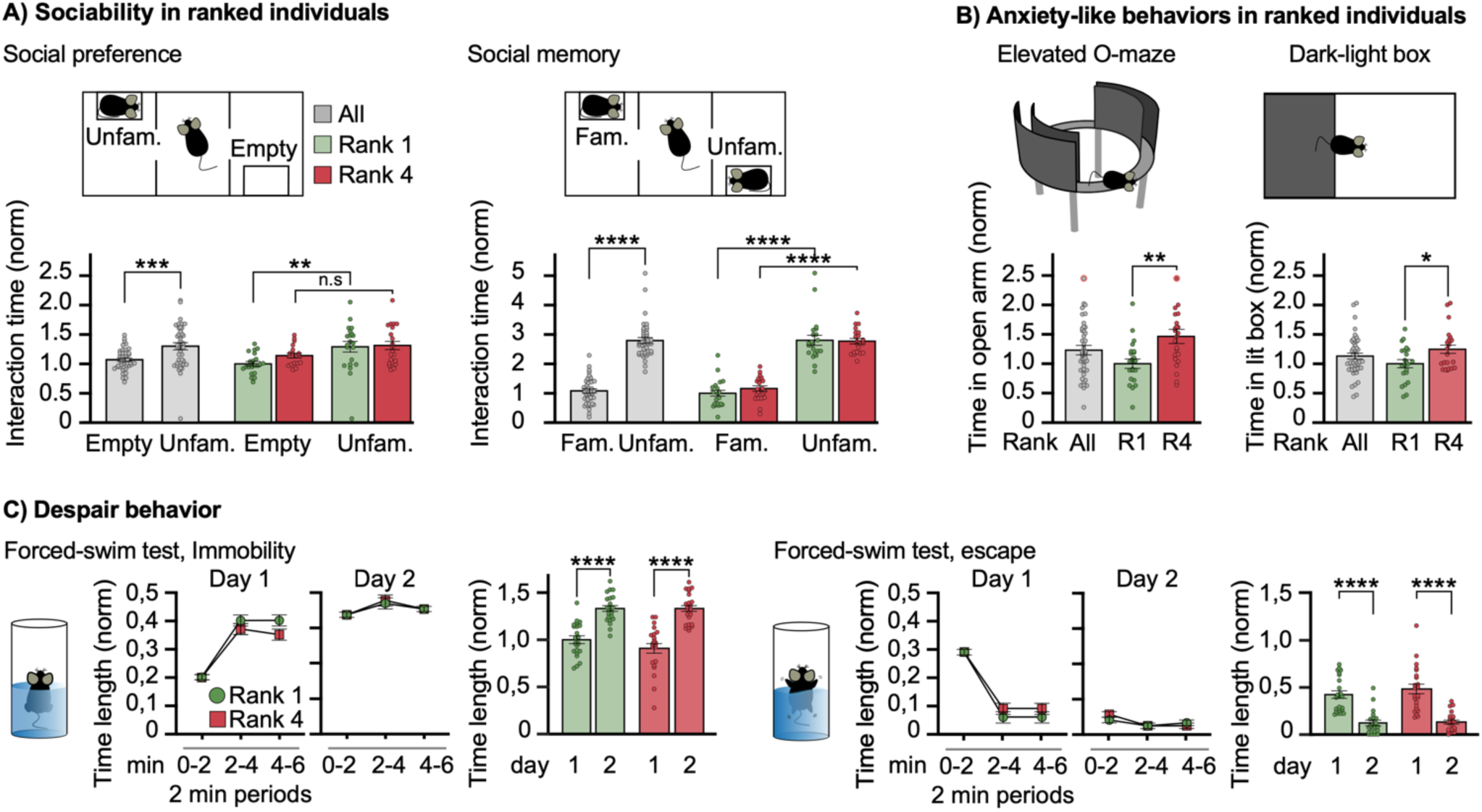
Behavioral differences are associated with social rank in genetically identical mice. **(A)** Higher-ranking mice show increased sociability but similar social memory compared to lower-ranking individuals. Left panel: social preference test (schematic shown above). Time spent interacting with an empty box (Empty) or a box containing an unfamiliar mouse (Unfam.) is shown for all C57BL/6 mice (grey, n = 42), including rank 1 (R1, green, n = 21), and rank 4 (R4, red, n = 21) individuals. Values were normalized to the mean interaction time of R1 mice with the empty box. Statistical analyses: all C57BL/6 mice, Wilcoxon matched-pairs signed-rank test. R1 *vs*. R4 mice: two-way repeated measures ANOVA followed by Bonferroni’s post hoc test. Right panel: social memory test. Time spent interacting with a familiar (Fam.) *vs.* an unfamiliar (Unfam.) mouse is shown for all C57BL/6 mice (grey, n = 42), including R1 (green, n = 21) and R4 (red, n = 21) individuals. Values were normalized to the mean interaction time of R1 mice with the familiar mouse. Statistical analyses: all mice, Wilcoxon matched-pairs signed-rank test; R1 *vs*. R4 mice: two-way repeated-measures ANOVA followed by Bonferroni’s post hoc test. ** P < 0.01, *** P < 0.001, **** P < 0.0001. Error bars represent ± SEM. **(B)** Anxiety-like behavior in ranked individuals. Left panel: elevated O-maze test. Time spent in the open sections is shown for all C57BL/6 mice (n = 45), including R1 (green, n = 22) and R4 (red, n *=* 23) individuals. Values were normalized to the mean time spent in the open sections by R1 mice. Statistical analysis: Mann Whitney test. One R4 data point outside the graph scale (3.36) is indicated by a red circle. Right Panel: dark-light box test. Time spent in the lit compartment is shown for all C57B/L6 mice (n = 44), including R1 (green, n = 22) and R4 (red, n = 22) individuals. Values were normalized to the mean time spent in lit compartment by R1 mice. Statistical analysis: Mann Whitney test. * P < 0.05, ** P < 0.01. Error bars represent ± SEM. **(C)** Despair-like behavior assessed using the forced swim test in R1 (n = 23) and R4 individuals (n = 23). Values were normalized to the mean immobility time of R1 mice on day 1. Left panel: time course of immobility duration measured in 2 min blocks on day 1 and day 2. Middle panel: total immobility duration. Right panel: time course of escape behavior duration course in 2-min blocks on day 1 and day 2. Statistical analysis: two-way repeated-measures ANOVA followed by Bonferroni’s post hoc test. **** P < 0.0001. Error bars represent ± SEM.

We next assessed social memory using the three-chamber test by comparing interaction times with a previously encountered (familiar) and a novel (unfamiliar) conspecific (Fig. 2A, top right). In contrast to social preference, social rank did not influence social memory. Both R1 and R4 individuals showed a marked preference for the unfamiliar conspecific (Fig. 2A, right panel). For all mice combined, interaction with the unfamiliar mouse was significantly greater than with the familiar mouse (P < 0.0001, Wilcoxon matched-pairs signed-rank test). Statistical analysis using two-way repeated-measures ANOVA followed by Bonferroni’s post hoc test revealed no interaction effect (P = 0.40; F(1,40) = 0.71) and no effect of social rank (P = 0.62; F(1,40) = 0.25), but a highly significant effect of familiarity (P < 0.0001; F(1,40) = 224.9). Post hoc comparisons confirmed that both R1 and R4 mice interacted significantly more with the unfamiliar than with the familiar conspecific (P < 0.0001 for both groups).

We next quantified anxiety-like behavior using two classical conflict-based exploration tests: the elevated O-maze and the dark-light tests (Fig. 2B, left and right, respectively). These assays rely on the innate aversion of mice to exposed, elevated or lit environments. R1 individuals spent significantly less time in the open sections of the O-maze than R4 individuals (n=22, n=23, green and red bars, respectively, Mann Whitney test P < 0.01). Similarly, the same R1 individuals spent also significantly less time in the illuminated compartment of the dark-light box than R4 individuals (Mann Whitney test P < 0.05). These findings indicate increased anxiety-like behavior in R1 mice, a pattern consistent with previous observations in males ^21 23^.

We also assessed despair-like behavior in the forced-swim test. Immobility and escape-related behaviors were quantified over a 6-minutes session repeated upon two consecutive days. No significant differences were observed between R1 and R4 individuals. On day 1, both groups showed a progressive increase in immobility over the course of the session (Fig. 2C, left panel; two-way repeated-measures ANOVA: no effect of social rank, P = 0.16, F(1,44) = 2.02; significant effect of time, P < 0.0001, F(2,88) = 103.0). As expected, immobility further increased on day 2 (two-way repeated-measures ANOVA: no effect of social rank, P = 0.92, F(1,44) = 0.01; significant effect of time, P < 0.001, F(2,88) = 8.08). Escape-related behavior showed the opposite temporal pattern (Fig. 2C, right panel).

Finally, we did not observe differences in general locomotor activity measured in a circular actometer between R1 and R4 mice (Fig. S4A). Analysis using two-way repeated-measures ANOVA followed by Bonferroni’s post hoc test revealed a significant interaction effect (P < 0.05; F(23,989) = 1.6) and a strong effect of time (P < 0.0001; F(23,989) = 59.0), but no overall effect of social rank (P = 0.19; F(1,43) = 1.8).

### Preexisting differences in sociability and possibly anxiety-like behaviors influence social fate

Behavioral differences observed between social ranks may either emerge as a consequence of adaptation to the social environment within the tetrad, or reflect pre-existing individual traits that predispose certain individuals to specific social outcomes, including rank attainment. To address this question, we singly housed C57BL/6 females for one week prior to tetrad formation and assessed their sociability and anxiety-like behaviors. Following tetrad formation and subsequent rank assignment, these pre-tetrad behavioral profiles were retrospectively stratified according to the individuals’ future social rank (Fig. 3A).

**Figure 3:**
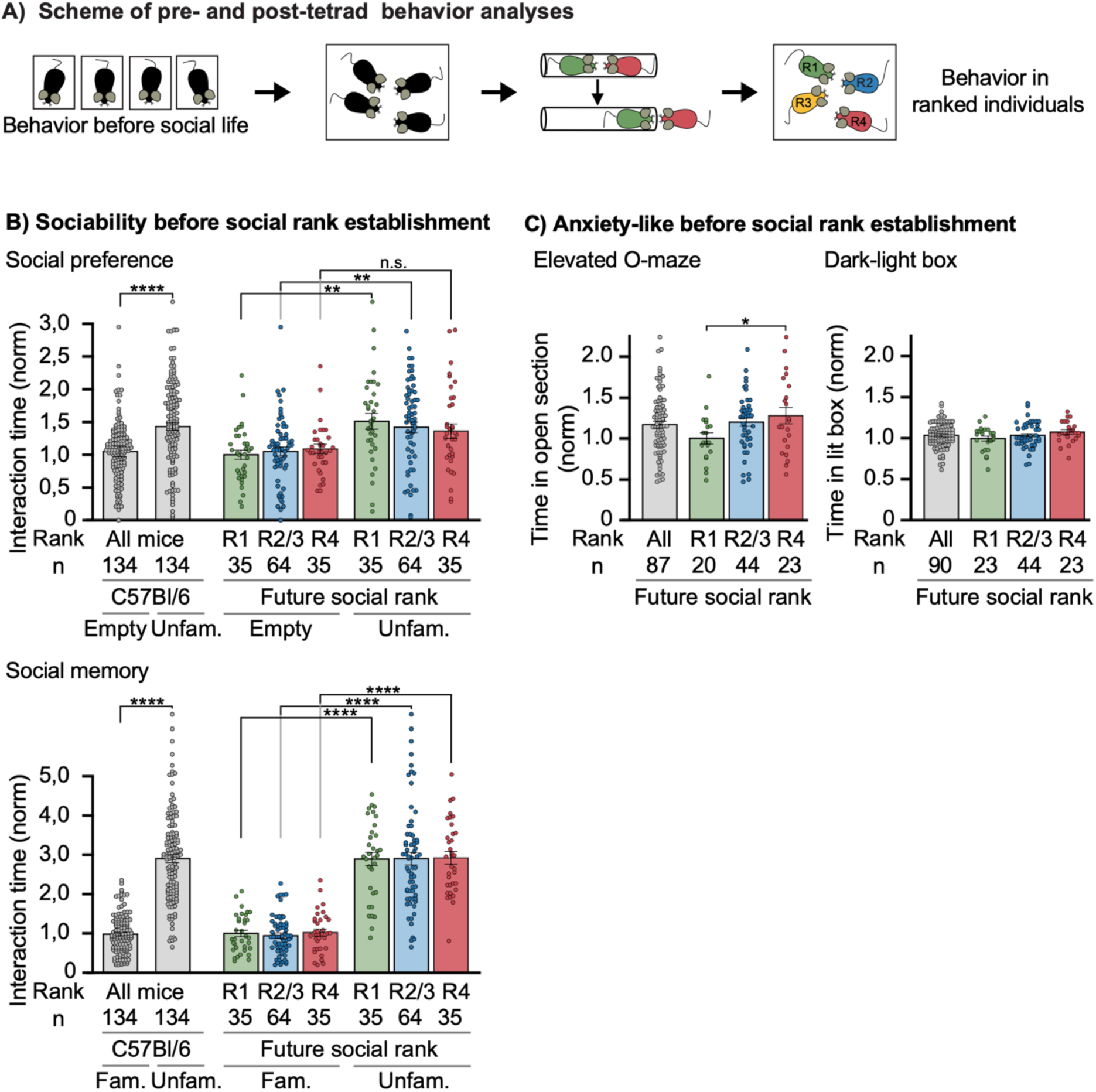
Behavioral differences between future rank 1 and rank 4 C57BL/6 female mice pre-exist the social hierarchy formation. **(A)** Experimental design. Prior to social housing, C57BL/6 female mice were singly housed and subjected to behavioral testing. **(B)** Social behavior before hierarchy formation. Top panel: social preference test. Future R4 individuals failed to display social preference, in contrast to future R1 mice. Shown is the time spent interacting with an unfamiliar conspecific (Unfam.) vs. an empty box (Empty) for all C57BL/6 mice from tetrads (grey bars, n= 134), including future R1 (green bars, n = 35), future R2/3 (blue bars, n = 64) and future R4 (red bars, n = 35) individuals. Values were normalized to the mean interaction time of future R1 mice with the empty box. Statistical analyses: all C57BL/6 female mice, Wilcoxon matched-pairs signed-rank test; ranked individuals, two-way repeated-measures ANOVA followed by Bonferroni’s post hoc test. Bottom panel: social memory test. Time spent interacting with a familiar (Fam.) *vs.* an unfamiliar (Unfam.) conspecific is shown. Values were normalized to the mean interaction time of future R1 mice with the familiar mouse. Statistical analyses: all C57BL/6 mice, Wilcoxon matched-pairs signed-rank test; ranked individuals, two-way repeated-measures ANOVA followed by Bonferroni’s post hoc test. ** P < 0.01, **** P < 0.0001. Error bars represent ± SEM. **(C)** Anxiety-like behavior before hierarchy formation. Values were normalized to the mean value of future R1 mice. Left panel: elevated O-maze test. Time spent in open-sections is shown for all C57BL/6 female mice (n = 87), including future rank 1 (green n = 20), future rank 2-3 (blue, n = 44) and future rank 4 (red, n = 23) individuals. Right panel: dark-light box test. Time spent in the lit compartment is shown. No significant differences in anxiety-like behavior were observed between future ranks (P = 0.23). Statistical analyses: Kruskal–Wallis test followed by Dunn’s post hoc test. * P < 0.05. Error bars represent ± SEM.

Future R1 (n = 35) and future R2/3 (n = 64) females displayed marked sociability whereas future R4 (n = 35) showed no significant preference for social interactions (Fig. 3B upper panel, green, blue and red bars, respectively). Statistical analysis using two-way repeated-measures ANOVA followed by Bonferroni’s post hoc test revealed no interaction effect (P = 0.55; F(2,131) = 0.60) and no effect of future social rank (P = 0.94; F(2,131) = 0.062), but a significant effect of social cue (P < 0.0001; F(1,131) = 21.04). Post hoc analysis showed significant social preference in future R1 and future R2/R3 mice (P < 0.01 for both groups), but not in future R4 individuals.

In contrast, social memory was similar across all groups, with all mice showing a consistent preference for the unfamiliar conspecific (Fig. 3B lower panel). Two-way repeated-measures ANOVA followed by Bonferroni’s post hoc test revealed no interaction effect (P = 0.98; F(2,135) = 0.021) and no effect of future social rank (P = 0.88; F(2,135) = 0.13), but a strong effect of familiarity (P < 0.0001; F(1,135) = 314.7). Post hoc comparisons confirmed significant preferences for the unfamiliar mouse in future R1, R2/R3, and R4 individuals (P < 0.0001 for all groups).

Anxiety-like behaviors showed a more complex pattern. In the elevated O-maze, future R1 individuals (n = 20) spent significantly less time in the open sections of the maze compared to future R4 ones (n = 23, Kruskal-Wallis test P < 0.05; Dunn’s post-hoc. P < 0.05), but did not differ significantly from future R2/R3 mice (n = 44; P = 0.058), suggesting increased anxiety-like behavior in future highest ranked individuals (Fig. 3C, left panel). In contrast, no such difference was observed in the dark–light test, where neither time spent in the illuminated compartment, nor the number of entries in the lit chamber differed significantly between groups (Fig. 3C, right panel, Kruskal–Wallis test, P = 0.23; and Fig. S5, Kruskal–Wallis test, P = 0.69).

Finally, we did not observe any differences in general locomotor activity between future R1 and R4 mice (Fig. S4B). Analysis using two-way repeated-measures ANOVA followed by Bonferroni’s post hoc test did not reveal an interaction effect (P = 0.62, F(46, 1587) = 0.93). There was a marked effect of time (P < 0.0001; F(23, 1587) = 144.9), but no overall effect of social rank (P = 0.76, F(2, 69) = 0.28).

Together, these findings indicate that social rank in female C57BL/6 mice is predicted by pre-existing sociability. The discrepant results obtained across anxiety-related paradigms do not allow us to draw a clear conclusion regarding the relationship between anxiety-like behavior and future social rank.

### In mixed-sex colonies, rank attainment is similar in males and females

To determine whether access to a specific social rank is sex-dependent and whether a social hierarchy emerges in a mixed-sex colony similarly to that in male- or female-only colonies, we generated tetrads composed of one male C57BL/6 mouse and three female C57BL/6 mice, and analyzed social organization using the precedence Tube-test. To avoid potential effects of body weight, male and female mice with similar body mass were grouped together (Fig. 4A, Fig. S6). In addition, to avoid behavioral confounds associated with gestation or lactation, male mice were vasectomized prior to colony formation. Two weeks later, we assessed social ranks using the Tube-test (session 1) and reassessed five weeks later (session 2). In the 12 mixed-sex tetrads analyzed male mice were evenly distributed across all ranks, indicating that sex does not seem to influence hierarchical position (Fig.4B). When ranks were reassessed one month later, 8 out of 12 R1 individuals and 7 out of 12 R4 individuals retained their original rank, suggesting that hierarchy stability may be lower than in male-only tetrads (Fig. 4C). As observed for male ^16 ,23^ and female tetrads, the extreme ranks in mixed colonies reached stability of the measurement in the Tube-test faster than intermediate ones (Fig. 4D). Half of the R1 individuals reached their final rank between days 1 and 2, and half of the R4 individuals between days 2 and 3, whereas half of R2 and R3 individuals stabilized between days 5 and 6 (Gehan-Breslow-Wilcoxon test R1 *vs.* R2: P < 0.05, R1 *vs*. R3: P < 0.05, R4 *vs*. R3: P < 0.05 and R4 *vs*. R2: P = 0.06). Focusing on contest dynamics, we observed that the male-female encounters during the Tube test lasted significantly longer than female-female encounters (5.9 ± 0.3 s, n = 324 *vs.* 4.8 ± 0.3 s, n = 324, respectively; unpaired t test, P < 0.05) (Fig. 4E). However, contest outcomes were not sex-biased. Males had a 57 ± 8 and 50 ± 10 % probability of winning during the first three days of the first and second Tube-test sessions, respectively; and 45 ± 10 and 41 ± 9 % probability during the last three days, when the measure was stabilized. A Wilcoxon Signed Rank Test indicated that none of these values differed significantly from chance level (50%) (Fig. 4F).

**Figure 4:**
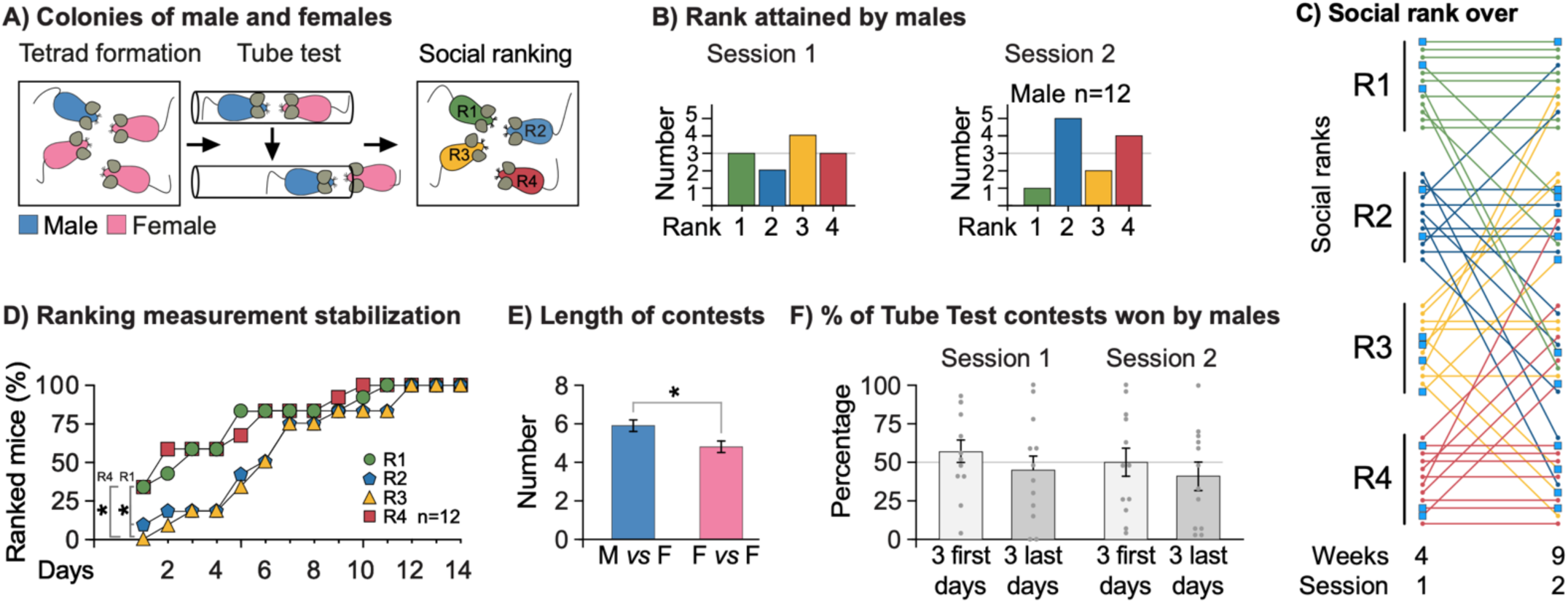
Social hierarchy analysis in mixed-sex tetrads. **(A)** Experimental design. Tetrads composed of one male and three female mice were generated and analyzed for social-rank establishment. **(B)** Distribution of social ranks among the 12 males from mixed-sex tetrads, assessed three weeks (session 1) and nine weeks (session 2) after tetrad formation. **(C)** Longitudinal evolution of male social ranks in 12 mixed-sex tetrads, between sessions 1 and 2. Ranks were assessed three and nine weeks after tetrad formation. Each row represents one mouse. Colors indicate the initial rank determined after three weeks: green (R1), blue (R2), orange (R3), and red (R4). Male individuals are indicated by a blue box. Within each rank category, the position of the dots indicates the tetrad to which the individual belonged, from Tetrad 1 (top) to Tetrad 12 (bottom). Changes in rank between sessions 1 and 2 are illustrated by connecting lines. **(D)** Stabilization dynamics of rank identification in the Tube-test. The cumulative percentage of individuals consistently assigned to each rank is shown across successive testing days (n = 12 tetrads, session 1). Gehan– Breslow–Wilcoxon test: * P < 0.05. Error bars represent ± SEM. **(E)** Duration of male–female and female–female contests during the Tube-test (n = 324 contests from 12 tetrads, session 1). Statistical analysis: Mann–Whitney test, * P < 0.05. **(F)** Percentage of contests won by males during the first three days and the last three days of tube testing, once stabilization of social organization measurement had been reached in sessions 1 and 2. Statistical analysis: Wilcoxon signed-rank test. Error bars represents ± SEM.

### Increased activity of dopamine neurons from the ventral tegmental area in highest-ranked females

We previously reported that R1 males exhibit reduced bursting activity of ventral tegmental area (VTA) dopamine neurons ^13^. We therefore investigated whether a similar difference could be observed by comparing VTA dopamine neuron activity between R1 and R4 female mice. To this end, we performed juxta-cellular single-unit recordings in anesthetized mice (Figure 5, left). Analysis of 39 neurons recorded from four R1 female and 40 neurons recorded from four R4 females unexpectedly revealed that both the spontaneous firing frequency and the percentage of spikes occurring within bursts was significantly higher in R1 individuals, (Wilcoxon signed-rank test, P < 0.001; and surrogate method, see Methods section, P < 0.05, respectively, Fig. 5, right graph). This pattern sharply contrasts with our previous observation in males, in which R1 mice displayed similar firing frequencies but reduced bursting activity compared to R4 males.

**Figure 5:**
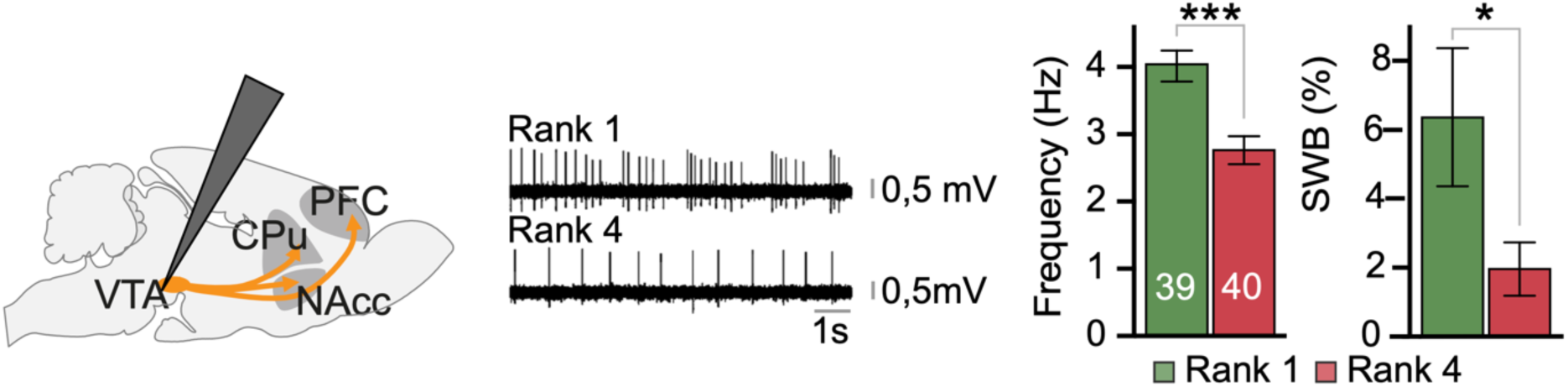
Dopamine neuron activity in the VTA is elevated in rank 1 females. Left panel: schematic representation of electrode placement for *in vivo* recordings. Middle panel: representative electrophysiological traces recorded from R1 and R4 mice. Right panel: quantification of the mean firing frequency (Hz) and percentage of spikes within bursts (%SWB) of VTA dopamine neurons recorded in R1 (n = 39 cells from 4 mice) and R4 (n = 40 cells from 4 mice) individuals. Statistical analyses: firing frequency, R1 *vs.* R4, *** P < 0.001; percentage of spikes within bursts, R1 *vs.* R4, * P < 0.05. Error bars represent ± SEM.

### In female mice, GR gene inactivation does not facilitate high rank attainment

In male mice, genetic ablation of the stress-responsive GR gene in dopaminoceptive neurons promotes access to the highest social rank ^21^. We therefore assessed whether the same applies to females by housing one adult GR^D1aCre^ female with three unfamiliar control (GR^loxP/loxP^) individuals (Figure 6, left panel), and evaluating social rank using the Tube-test two weeks later.

**Figure 6:**
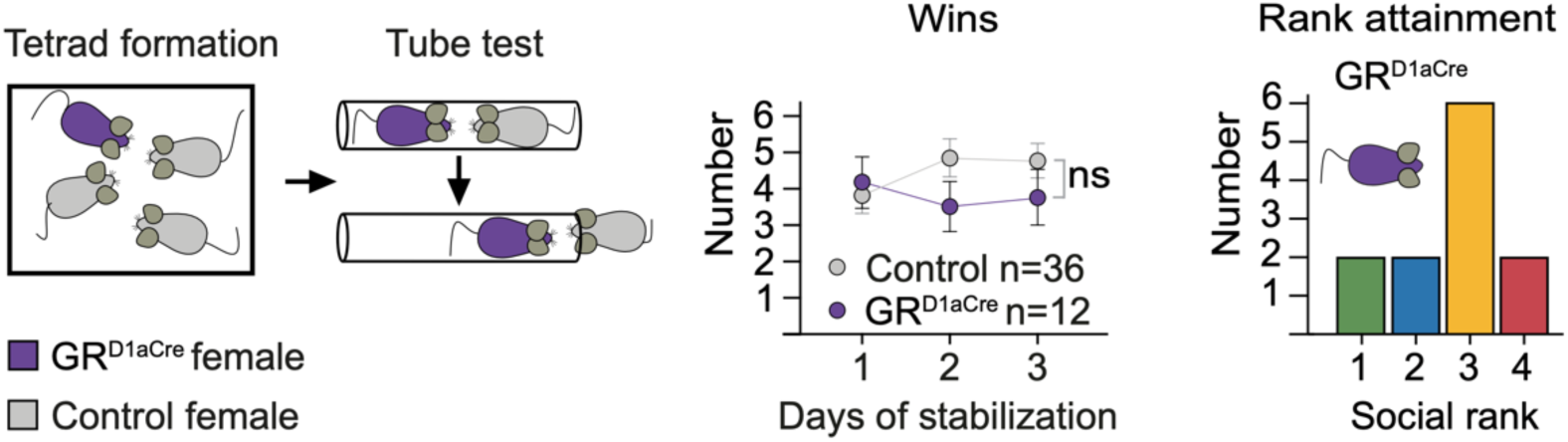
Disruption of GR signaling in dopaminoceptive neurons does not alter social-rank attainment in female mice. Left panel: schematic representation of tetrads composed of one GR^D1aCre^ mutant female and three control females, and corresponding rank assignment following Tube-test assessment. Middle panel: cumulative number of wins during the last three days of stabilization of the Tube-test for control mice (grey) and GR^D1aCre^ mutant (purple) females. Statistical analysis: two-way repeated-measures ANOVA followed by Bonferroni’s post hoc test. Error bars represent ± SEM. Right panel: distribution of social ranks attained by GR^D1aCre^ females across 12 tetrads.

Unlike in male groups, GR^D1aCre^ females did not exhibit a significantly higher average number of daily wins during the last three days of the Tube-test compared to controls (Fig. 6, middle chart). A significant interaction between genotype and days even indicates that the number of wins in GR^D1aCre^ females rather decreases across the last 3 days of stabilization (two-way repeated measured ANOVA with Bonferroni’s post hoc test: interaction effect, P < 0.0001, F(2, 92) = 13.23; effect of time, P < 0.05, F(2, 92) = 3.308; no effect of genotype, P = 0.4773, F(1, 46) = 0.5133). Accordingly, GR gene inactivation did not confer any advantage in attaining a social rank in female mice.

## Discussion

The mechanisms underlying the emergence of social organization in mice have mostly been studied in male colonies. In the present study, we sought to determine whether the mechanisms shaping male social hierarchy also apply to females.

As reported by several authors, we observed that female mouse colonies establish stable social hierarchies that can be assessed through a precedence-based test (Tube-test) or resource competition (e.g., food or warmth) ^25–33^. Female hierarchies were however less stable than male ones over time.

Similar to males, rank 1 female displayed increased anxiety-like behavior and sociability compared with rank 4 females ^21, 23^. This is consistent with previous finding in paired C57BL/6 females, in which highest ranked individuals displayed increased sociability and anxiety ^30^. Another study using the the same mouse strain but grouped in colonies of five individuals, did not observe any anxiety differences between ranks in either males or females ^27^.

Behavioral correlates of social rank in established groups have been studied in many vertebrate species but only a limited number of studies have identified individual traits that predict the social fate of individuals. Greater boldness in bold trout, shorter stress responses in rainbow trout, and stronger exploratory behavior in birds such as mountain chickadees and great tits have all been associated with a higher probability of attaining elevated social status ^37^. By profiling individual behavior before the formation of female tetrads, we observed that future R1 female were more prone to engage in social interactions, similar to what we previously observed in male ^21^. Regarding anxiety-like behaviors, we did not observe differences between future rank 1 and rank 4 males, whereas results in females were less consistent, with increased anxiety detected in the elevated O-maze but not in the dark-light box test. Thus, as in males, pre-existing sociability appears to shape social status trajectories within a female colony. In C57B/L6 females, Smith-Osborne *et al.* ^30^ reported increased exploratory rearing in future dominant individuals prior to dyad-housing. Excluding genetic differences when using isogenic individuals, we assume that prior behavioral differences arise from previous life experiences. In mice, prenatal factors, such as maternal stress exposure or as well as postnatal conditions including maternal care, can shape adult behavior ^38–41^. Adolescence also represents a critical developmental period, as illustrated by the long-term effects of social isolation on adult anxiety and sociability ^42–44^. Isolation has prominent anxiogenic effect in male rats when it occurs shortly after weaning but an anxiolytic effect if it occurs after day 45. In female, rats, subtle or no effects have been reported ^42, 43^. Many studies compare animals isolated during juvenile stages with animals isolated only during adulthood, without resocialization before behavioral testing. When examined, resocialization after late isolation frequently reverses the behavioral effects ^45, 42, 43^. In our experiments, seven-week-old females were singly housed for 10 days before tetrad formation in order to perform behavioral profiling. We therefore cannot exclude the possibility that this procedure exacerbated behavioral differences between future rank 1 and rank 4 individuals.

The simplicity of the Tube-test to identify a stable social organization within a group of mice made its recent success. Whether this social organization reflects what has been classically described as “social dominance” usually measured by aggressivity, territoriality and access to resources raised questioning for the last fifty years ^46^. In male mice, aggressivity is a key component of the definition of social hierarchy and, we previously observed the Tube-test defined C57BL/6 rank 1 individual are less aggressive than rank 4 ones. Comparing four classical tests of “dominance” including agonistic behavior, urine marking, Tube-test and access to a resource, in C57BL/6 and CD1 mouse males, Cum *et al.* ^36^ observed a similar inverted correlation between agonistic behavior and test tube rankings for CD1 mice and an absence of correlation for C57BL/6 males. In C57BL/6 males, antagonistic behavior and urine marking were positively correlated, as were Tube-test and resource access but with the two first inversely to the two last. In C57BL/6 mice access to a resource, such a warm plate in a cold room allowed us to identify lowest ranked females in the Tube-test, and highest ranked males, similar to other observations ^33^. Interestingly, in a semi-naturalistic setting, Reinwald *et al.* ^47^ used an automated Tube-test and observed a stable social organization with this criterium, that was correlated to an antagonistic behavior (chasing), the correlation being stronger in groups with less clearly defined hierarchies. Altogether, these data may suggest that social organization of a group is multidimensional resulting from a dynamic interplay between distinct but overlapping hierarchies, either influenced by strength, aggressivity, precedence, social experience, *etc*.

This organization must be complexified in colonies formed of males and females. Considering social rank and sex, it is generally assumed that males dominate females, particularly in species displaying male-biased sexual dimorphism in body size and weight. Ethological observations across species, however, suggest that the situation is often more nuanced. ^48^.

Given the behavioral similarities observed between highest- and lowest-ranked individuals in same-sex male and female C57BL/6 colonies, both before and after colony formation, we asked whether mixed-sex groups could also organize hierarchically and whether social rank attainment would depend on sex. In mixed-sex tetrads, composed of three females and one male, we observed a clear ranking of the individuals, though the social organization seems less stable over time than in only-male colonies. Males and females showed a similar probability of winning Tube-tests contests, although male-female interactions were significantly shorter than female-female interactions, suggesting that sex affects encounter dynamics. Importantly, no sex-bias in rank attainment was detected. These findings resemble observations in mixed-sex rat colonies housed in a large semi-naturalistic environment (Visible Burrow System), where females, despite being substantially smaller than males, dominated approximately 55% of males on average, and in several colonies a female attained the highest social rank ^49^.

In males, the mesocorticolimbic dopamine system has been implicated in social rank regulation. D1-receptor antagonism promotes dominance in mid-ranked mice, whereas D2-receptor antagonism decreases rank in both mice and macaques ^50, 51^. In the prefrontal cortex, D2-receptor inactivation lowers social ranking, while ex vivo studies revealed increased D1 receptor activity in top-ranked mice and elevated D2 receptor activity in low-ranked individuals ^52^. In the nucleus accumbens, D1 receptor activation is associated with social contests, and its blockade reduces dominance behavior ^53^.

We previously found in males that spontaneous VTA dopamine neuron activity, measured by extracellular recordings in anaesthetized animals, is reduced in top-ranked individuals ^21^. This basal firing rate likely reflects an intrinsic, stable property of the dopaminergic system, one that is insulated from the immediate sensory and behavioral context, and may therefore constitute a meaningful correlate of a lasting trait such as social rank, rather than a transient response to social interaction. A key observation that helps establish the functional relevance of VTA dopaminergic activity to hierarchy formation in males is that experimentally inhibiting VTA dopaminergic neurons was sufficient to promote access to a higher social rank. This suggests that in males, dopaminergic tone is not merely a read-out of an already-established hierarchy but rather an active participant in its emergence. Converging evidence came from the observation that selective ablation of glucocorticoid receptors in dopamine-sensitive neurons, which reduces stress responsiveness, similarly decreases spontaneous VTA dopaminergic firing in anesthetized animals and promotes higher rank attainment ^34, 21^.

In contrast to males, rank 1 female displayed significantly higher VTA dopamine neuron firing rates and bursting activity than rank 4 females. Sex-dimorphism of the VTA dopaminergic system that modulate different behaviours in males and females have been reported in several contexts. For example, higher-ranked males are resilient to psychostimulant-seeking behavior, whereas lower-ranked males and females are not, potentially due to reduced nucleus accumbens dopaminergic function specifically in high-ranked males ^54^. Another recent study reported sex differences in spontaneous VTA neuronal activity in triads, in which males formed stable worker-scrounger relationships driven by competition, whereas female triads adopted more uniform and cooperative strategies ^55^. Sex-based differences in VTA activity are also reported in stress paradigms, including opposite effects of oxytocin signaling on defeat-induced social avoidance in males and females ^56^, and sex-dependent modulation of VTA firing rates following sub-chronic stress ^57^. Interestingly, Wei *et al*. ^58^ showed that sexually dimorphic modes of dopaminergic encode the regulation of context-dependent social preferences, supporting the idea that also similar social outcomes can emerge from distinct dopamine-dependent processes in males and females. Estrogen may contribute to this modulation of VTA excitability ^59, 60^. A limitation of our study is the fact that we did not monitor estrous cycle stages. However, studies in female rat triads and mouse groups reported that estrous cycle phase does not significantly influence social rank ^61, 62^. However conversely, social rank itself may affect estrous cyclicity since dominant females have been reported to exhibit longer estrous cycles. Williamson *et al.* (2019) ^28^ showed that if social rank influences estrous cyclicity in CD1 dominant female, estrous cycle does not affect social hierarchy based on agonistic interactions. In the same mouse strain, Rieneker *et al.* (2019) ^61^ observed also an absence of estrus effects, this time, on Tube-test outcomes when a female in estrus faced a conspecific that was not. Similarly, in female rats, estrus did not shape social ranking based on play behavior ^62^.

The opposite relationship between social rank and spontaneous VTA activity observed in males and females suggests that VTA dopaminergic tone may not encode dominance itself, but rather distinct socioemotional dispositions whose nature, e.g. stress resilience in males, or social motivation in females, maps onto sex-specific processes that participate to the establishment of social ranking. This question is particularly interesting from an evolutionary point of view to understand the male-female and intra-sex powers relationships on resource-holding and mating since asymmetries on both shaping the social organizations and the mating systems of mammalian societies ^63^. If the role of coercive social control or dominance on social organization have been largely studied, knowledge over physiological mechanisms underlying other key aspects, including sharing, cooperation, interindividual bounding, *etc*. is more elusive, in particular in the context of mixed-sex colonies. These different aspects will be essential to better understand the dynamics of social organizations .and their diversity through evolution.

Finally, in many vertebrate species, including humans, stress-induced glucocorticoid levels are affected by social ranking ^29, 64–70^. Most studies have focused on males, and the few examining female rodents report either elevated levels in lower ranked female mice ^28^, or no association ^30^, although in this later case future ranking was predicted by glucocorticoid levels. In male mice, we previously showed that GR gene inactivation in dopaminoceptive neurons promotes high rank ^18, 21^. This effect was not reproduced in females, suggesting that if stress-elicited glucocorticoid release modulates social organization in female colonies, they do not act through this neuronal population.

Taken together, our results demonstrate that social organization emerges in both female and male colonies of genetically identical mice. Pre-existing individual traits influence social fate in both sexes but female hierarchies appear more fluid and governed by distinct physiological mechanisms.

## Acknowledgements

The authors thank Laurence Amar, Dorian Battivelli, and members of the GRAB team for helpful discussions and critical reading of the manuscript. The authors wish to thank Fabrice Machulka, Jean Vincent and other members of the IBPS animal facilities. This work was supported by the Foundation for Medical Research (FRM-Equipe grant DEQ20140329552 to FT), the Labex BioPsy, the foundation for brain research (FRC Neurodon, SP), the INCA (TABAC-19_020 grant to PF and FT). A preprint version of this work has been deposited on BioRXiv and HAL databases.

## Contributions

F.T. and S.P conceived and designed the experiments. E.C., C.V., A.G., F.M, P.F., S.P. and F.T. acquired and analyzed the data. E.C., S.P. and F.T. wrote the manuscript, and all authors revised it.

## Declaration of AI-assisted technologies in the process

During the preparation of the manuscript, the authors used ChatGPT in order to correct grammatical errors and improve English language. After using this tool, the authors reviewed and edited the content as needed and take full responsibility for the content of the publication.

## Supplemental information

### Social hierarchy and behavioral individuality in colonies of isogenic female mice

**Figure S1.**
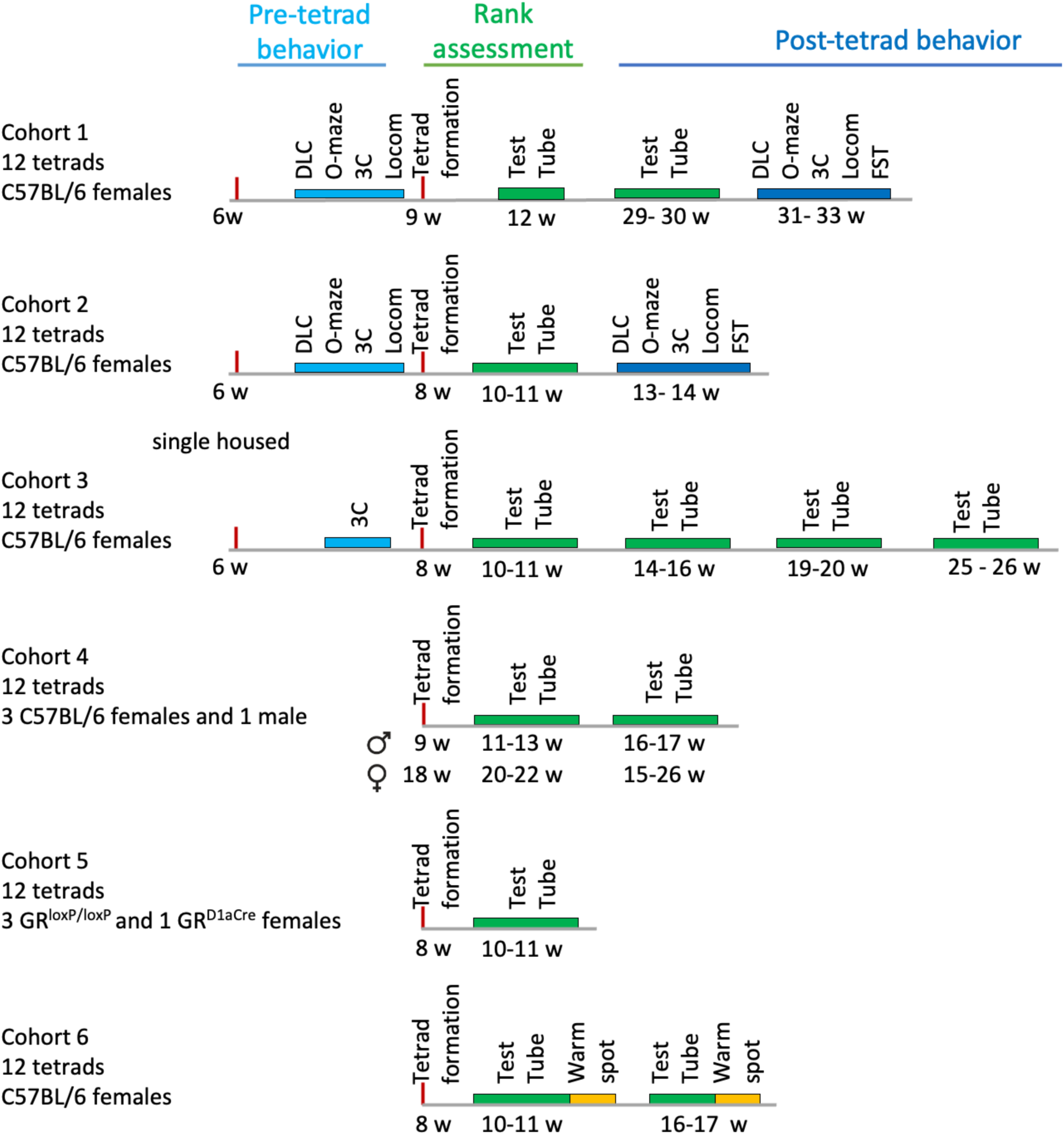
Experimental timeline for studies using ranked tetrads. Schematic representation of the sequential experimental procedures conducted before and after the establishment of social organization in tetrads. Ranking procedure are indicated by green and orange boxes (Tube-test and Warm-spot test, respectively), whereas blue boxes denote behavioral assays: DLB (dark-light Box), O-Maze (elevated O-maze), 3C (three-chamber sociability test), FST (forced-swim test), and Locom (locomotor activity). The ages of the mice at each timepoint is indicated below the corresponding boxes in weeks (w).

**Figure S2.**
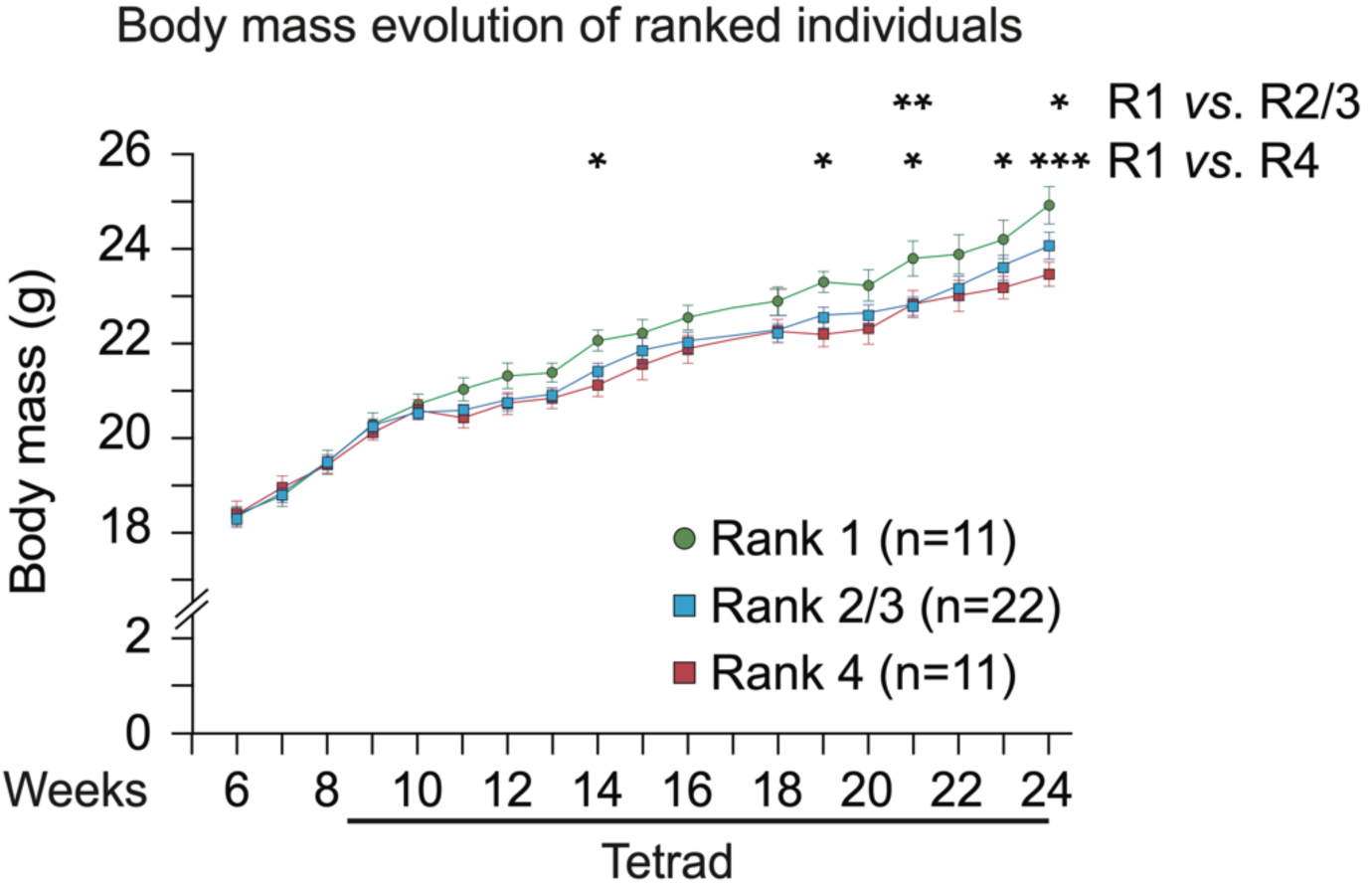
Longitudinal changes in body mass according to social rank determined by the initial Tube-test. Average body mass of individuals from each rank measured over a 16-week period. Time spent in tetrads is indicated below by the black line. Repeated two-way ANOVA followed by Bonferroni’s post hoc test, * P < 0.05, ** P < 0.01, *** P < 0.001. Error bars represent ± SEM

**Figure S3.**
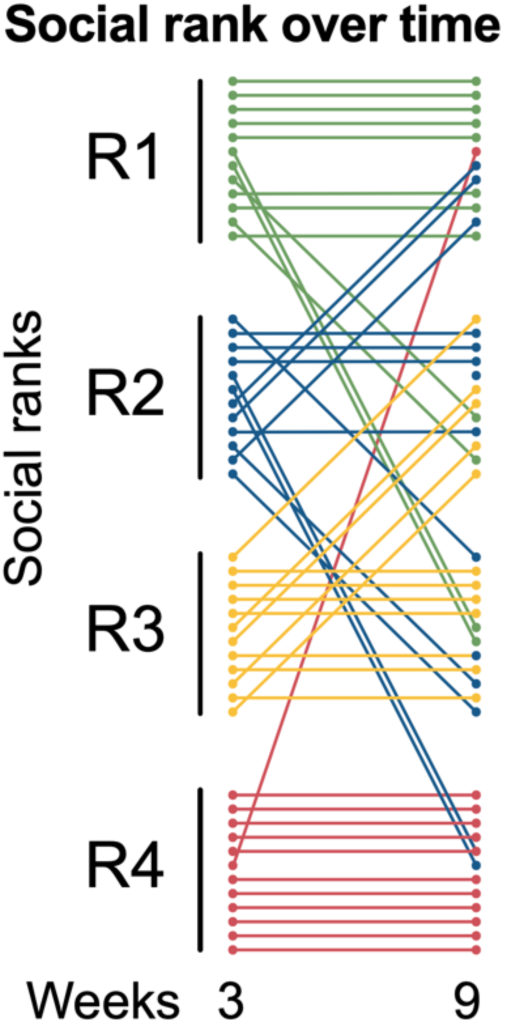
Repeated social rank assessment by Tube-test in the C57BL/6 female tetrads analyzed in the Warm spot assay. Twelve tetrads were assessed twice, three and nine weeks after tetrad formation. Each row represents one mouse. Colors indicate the initial rank determined after three weeks: green (R1), blue (R2), orange (R3), and red (R4). At the three-week time point, the position of the dots within the R1, R2, R3 and R4 groups indicates the tetrad to which the individual belonged, from Tetrad 1 (top) to Tetrad 12 (bottom). Changes in rank over time are illustrated by connecting lines.

**Figure S4.**
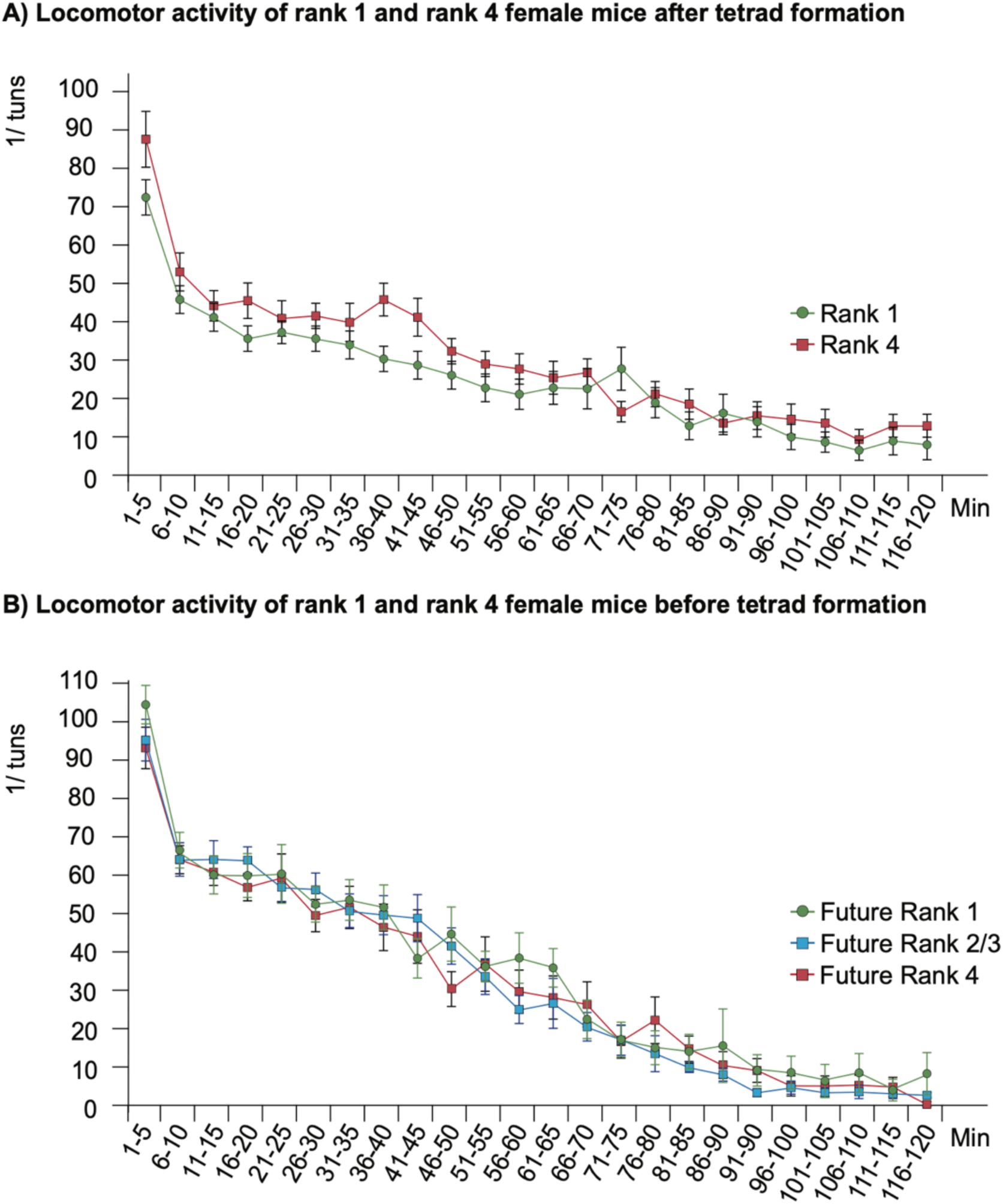
Basal locomotor activity in female mice after and before social housing. **(A)** Time course of locomotor activity in ranked individuals, measured as quarter turns in 5 min intervals. (R1, green, n = 22; R4, red, n = 23). **(B)** Time course of locomotor activity measured before tetrad formation in future R1 (n = 19), future R2/R3 (blue, n = 33), and future R4 (n = 20) individuals. Two-way repeated-measures ANOVA followed by Bonferroni’s post hoc test. No effect of rank was observed. Error bars represent ± SEM.

**Fig. S5.**
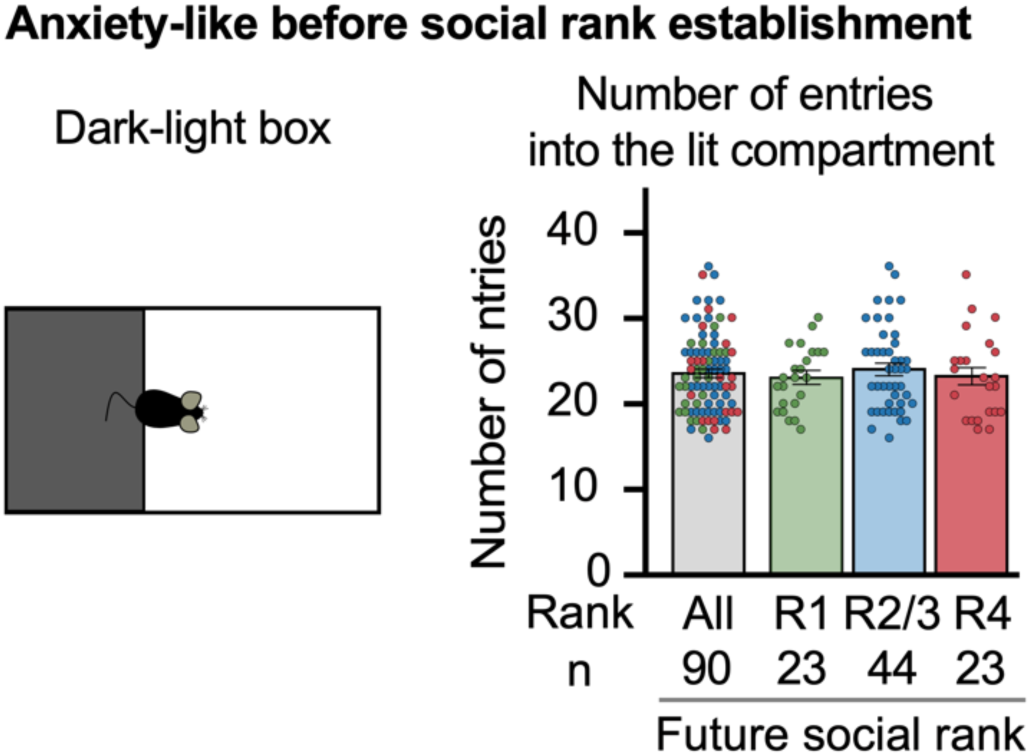
Number of entries in the lit compartment in the dark-light box test before tetrad formation. The future social ranks of C57BL/6 female mice, as well as the n are indicated below the graph. Kruskal–Wallis test. Error bars represent ± SEM.

**Figure S6.**
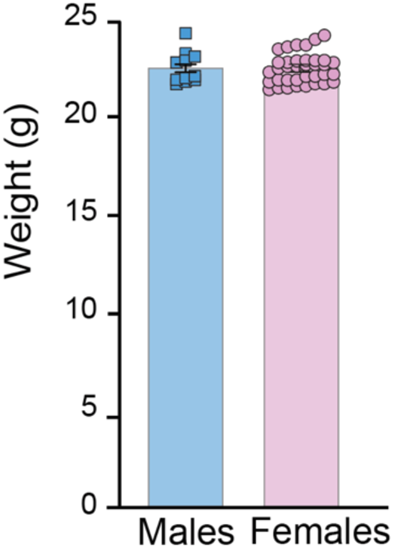
Body mass of male and female mice at the time of mixed tetrad formation. Body mass of males (blue, n = 12) and females (pink, n = 36) at the time of mixed-sex tetrad formation. Mann–Whitney nonparametric test, P = 0.96. Error bars represent ± SEM.

## References

1. Tinbergen, N. (1939). On the analysis of social organization among vertebrates, with special reference to birds. Am. Midl. Nat. 21, 210. 10.2307/2420381.

2. Francis, R.C. (1984). The effects of bidirectional selection for social dominance on agonistic behavior and sex ratios in the paradise fish (Macropodus Opercularis). Behaviour 90, 25–44. 10.1163/156853984X00542.

3. Chambers, L.K., Singleton, G.R., and Krebs, C.J. (2000). Movements and social organization of wild house mice (*Mus domesticus*) in the wheatlands of northwestern Victoria, Australia. Journal of Mammalogy 81(1): 59–69. 10.1644/1545-1542(2000)081<0059:MASOOW>2.0.CO;2

4. Hurst, J.L. (1987). Behavioural variation in wild house mice Mus domesticus Rutty: A quantitative assessment of female social organization. Animal Behaviour 35, 1846–1857. 10.1016/S0003-3472(87)80077-5.

5. Bronson, F.H. (1979). The reproductive ecology of the house mouse. Q Rev Biol 54, 265–299. 10.1086/411295.

6. Rolland, C., MacDonald, D.W., Fraipont, M.D., and Berdoy, M. (2003). Free Female Choice in House Mice: Leaving Best for Last. Behaviour 140, 1371–1388. 10.1163/156853903771980639.

7 Wang, F., Kessels, H.W., and Hu, H. (2014). The mouse that roared: neural mechanisms of social hierarchy. Trends Neurosci. 37(11):674–82. 10.1016/j.tins.2014.07.005.

8. Zheng, H., Chen, D., Zhong, Z., Li, Z., Yuan, M., Zhang, Z., Zhou, X., Zhu, G., Sun, H., and Sun, L. (2025). Behavioral tests for the assessment of social hierarchy in mice. Front Behav Neurosci. 19:1549666. 10.3389/fnbeh.2025.1549666.

9. Blanchard, D.C., Fukunaga-Stinson, C., Takahashi, L.K., Flannelly, K.J., and Blanchard, R.J. (1984). Dominance and aggression in social groups of male and female rats. Behav Processes. 9(1):31–48. 10.1016/0376-6357(84)90006-8.

10. Blanchard, R.J., Dulloog, L., Markham, C., Nishimura, O., Nikulina Compton, J., Jun, A., Han, C., and Blanchard, D.C. (2001). Sexual and aggressive interactions in a visible burrow system with provisioned burrows. Physiol Behav. 72(1-2):245–54. 10.1016/s0031-9384(00)00403-0.

11. Zilkha, N., Chuartzman, S.G., Sofer, Y., Pen, Y., Cum, M., Mayo, A., Alon, U., and Kimchi, T. (2023). Sex-dependent control of pheromones on social organization within groups of wild house mice. Curr Biol. 33(8):1407–1420. 10.1016/j.cub.2023.02.039.

12. Crowcroft, P., and Rowe, F.P. (1963). Social organization and territorial behaviour in the wild house mouse (Mus musculus L.). Proc. Zool. Soc. Lond. 140:517–531. 10.1111/j.1469-7998.1963.tb01871.

13. Vogt CC, Zipple MN, Sprockett DD, Miller CH, Hardy SX, Arthur MK, Greenstein AM, Colvin MS, Michel LM, Moeller AH, Sheehan MJ. (2024). Female behavior drives the formation of distinct social structures in C57BL/6J versus wild-derived outbred mice in field enclosures. BMC Biol. 22(1):35. 10.1186/s12915-024-01809-0.

14. Zipple MN, Chang Kuo D, Meng X, Reichard TM, Guess K, Vogt CC, Moeller AH, Sheehan MJ. (2025). Competitive social feedback amplifies the role of early life contingency in male mice. Science. 387(6729):81–85. 10.1126/science.adq0579.

15. Hollis, F., van der Kooij, M.A., Zanoletti, O., Lozano, L., Cantó, C., and Sandi, C. (2015). Mitochondrial function in the brain links anxiety with social subordination. P.N.A.S. 112, 15486–15491. 10.1073/pnas.1512653112.

16. Wang, F., Zhu, J., Zhu, H., Zhang, Q., Lin, Z., and Hu, H. (2011). Bidirectional Control of Social Hierarchy by Synaptic Efficacy in Medial Prefrontal Cortex. Science. 334, 693–697. 10.1126/science.1209951.

17. Cordero, M.I., and Sandi, C. (2007). Stress amplifies memory for social hierarchy. Front Neurosci. 1(1):175–84. 10.3389/neuro.01.1.1.013.2007.

18. Costa, D.F., Moita, M.A., and Márquez, C. (2021). Novel competition test for food rewards reveals stable dominance status in adult male rats. Sci Rep. 11(1):14599. . 10.1038/s41598-021-93818-0.

19. Padilla-Coreano, N., Batra, K., Patarino, M., Chen, Z., Rock, R.R., Zhang, R., Hausmann, S.B., Weddington, J.C., Patel, R., Zhang, Y.E., Fang, H.S., Mishra, S., LeDuke, D.O., Revanna, J., Li, H., Borio, M., Pamintuan, R., Bal, A., Keyes, L.R., Libster, A., Wichmann, R., Mills, F., Taschbach, F. H., Matthews, G.A., Curley, J.P., Fiete, I.R., Lu, C., and Tye, K.M. (2022). Cortical ensembles orchestrate social competition through hypothalamic outputs. Nature, 603(7902), 667–671. 10.1038/s41586-022-04507-5.

20. Zhou. T., Zhu, H., Fan, Z., Wang, F., Chen, Y., Liang, H., Yang, Z., Zhang, L., Lin, L., Zhan, Y., Wang, Z., and Hu, H.. (2017). History of winning remodels thalamo-PFC circuit to reinforce social dominance. Science 357: 162–168. 10.1126/science.aak9726.

21. Battivelli, D., Vernochet, C., Conabady, E., Nguyen, C., Zayed, A., Lebel, A., Meirsman, A.C., Messaoudene, S., Fieggen, A., Dreux, G., Rigoni, D., Le Borgne, T., Marti, F., Contesse, T., Barik, J., Tassin, J.P., Faure, P., Parnaudeau, S., and Tronche, F. (2024). Dopamine neurons activity and stress signalling as links between social hierarchy and vulnerability to psychopathologies. Biological Psychiatry. 95(8):774–784. 10.1016/j.biopsych.2023.08.029.

22. Lindzey, G., Winston, H., and Manosevitz, M. (1961). Social dominance in inbred mouse strains. Nature. 191:474–6. 10.1038/191474a0.

23. Larrieu, T., Cherix, A., Duque, A., Rodrigues, J., Lei, H., Gruetter, R., and Sandi, C. (2017). Hierarchical status predicts behavioral vulnerability and nucleus accumbens metabolic profile following chronic social defeat stress. Current Biology. 27, 2202–2210. 10.1016/j.cub.2017.06.027.

24. Papilloud, A., Weger, M., Bacq, A., Zalachoras, I., Hollis, F., Larrieu, T., Battivelli, D., Grosse, J., Zanoletti, O., Parnaudeau, S., Tronche, F., and Sandi, C. (2020). The glucocorticoid receptor in the nucleus accumbens plays a crucial role in social rank attainment in rodents. Psychoneuroendocrinology. 112, 104538. 10.1016/j.psyneuen.2019.104538.

25. Schuhr, B. (1987) Social structure and plasma corticosterone level in female albino mice. Physiol Behav. 40(6):689–93. 10.1016/0031-9384(87)90269-1.

26. van den Berg, W.E., Lamballais, S., Kushner, S.A. (2015) Sex-specific mechanism of social hierarchy in mice. Neuropsychopharmacology. 40(6):1364–72. 10.1038/npp.2014.319.

27. Varholick, J.A., Bailoo, J.D., Palme, R., and Würbel, H. (2018). Phenotypic variability between Social Dominance Ranks in laboratory mice. Scientific Report. 8, 6593. 10.1038/s41598-018-24624-4.

28. Williamson, C.M., Lee, W., DeCasien, A.R., Lanham, A., Romeo, R.D., and Curley, J.P. (2019). Social hierarchy position in female mice is associated with plasma corticosterone levels and hypothalamic gene expression. Scientific Reports. 9, 7324. 10.1038/s41598-019-43747-w.

29. LeClair, K.B., Chan, K.L., Kaster, M.P., Parise, L.F., Burnett, C.J., and Russo, S.J. (2021). Individual history of winning and hierarchy landscape influence stress susceptibility in mice. Elife. 10: e71401. 10.7554/eLife.71401.

30. Smith-Osborne, L., Duong, A., Resendez, A., Palme, R., and Fadok J.P. (2023). Female dominance hierarchies influence responses to psychosocial stressors. Current biology. 33(8), 1535–1549.e5. 10.1016/j.cub.2023.03.020.

31. Fulenwider, H.D., Zhang, Y., and Ryabinin, A.E. (2024). Characterization of social hierarchy formation and maintenance in same-sex, group-housed male and female C57BL/6 J mice. Hormones and behavior, 157, 105452. 10.1016/j.yhbeh.2023.105452.

32. Sofer, Y., Zilkha, N., Gimpel, E., Wagner, S., Chuartzman, S.G., and Kimchi, T. (2024). Sexually dimorphic oxytocin circuits drive intragroup social conflict and aggression in wild house mice. Nat Neurosci. 27(8):1565–1573. 10.1038/s41593-024-01685-5.

33. Zheng, D., Xin, Q., Jin, S., Zhou, A., Jia, X., Tan, Y., and Hu, H. (2025). Neural mechanism of the sexually dimorphic winner effect in mice. Neuron. 113(24):4217–4231.e6. 10.1016/j.neuron.2025.09.029.

34. Ambroggi, F., Turiault, M., Milet, A., Deroche-Gamonet, V., Parnaudeau, S., Balado, E., Barik, J., van der Veen, R., Maroteaux, G., Lemberger, T., Schütz, G., Lazar, M., Marinelli, M., Piazza, P.V., and Tronche, F. (2009). Stress and addiction: glucocorticoid receptor in dopaminoceptive neurons facilitates cocaine seeking. Nature Neuroscience 12, 247–49. 10.1038/nn.2282.

35. Grace, A., and Bunney, B. (1984). The control of firing pattern in nigral dopamine neurons: burst firing. Journal of Neuroscience. 4, 2877–2890. 10.1523/JNEUROSCI.04-11-02877.1984.

36. Cum, M, Santiago Pérez, J.A., Iwata, R.L., Lopez, N., Higgs, A., Li, A., Ye, C., Wangia, E., Wright, E.S., García Restrepo, C., and Padilla-Coreano, N. (2024). A Multiparadigm Approach to Characterize Dominance Behaviors in CD1 and C57BL6 Male Mice. eNeuro. 11(11):ENEURO.0342-24.2024. 10.1523/ENEURO.0342-24.2024.

37. Chichinadze, K., Chichinadze, N., Gachechiladze, L., Lazarashvili, A., and Nikolaishvili, M. (2014). Physical predictors, behavioural/emotional attributes and neurochemical determinants of dominant behaviour. Biol Rev Camb Philos Soc. 89(4):1005–20. 10.1111/brv.12091.

38. Lee, Y.A., Kim, Y.J., and Goto, Y. (2016). Cognitive and affective alterations by prenatal and postnatal stress interaction. Physiol Behav. 15;165:146–53. 10.1016/j.physbeh.2016.07.014.

39. Curley, J.P., Champagne, F.A. Shaping the development of complex social behavior. (2023). Ann N Y Acad Sci. 1530(1):46–63. 10.1111/nyas.15076.

40. Francis-Oliveira, J., Tanaka, R., Shen, M., Cruvinel, E., Kano, S.I., and Niwa, M. (2026). Adolescent social isolation creates a latent vulnerability in maternal care with intergenerational social consequences, rescued by experienced mothers. bioRxiv [Preprint] 10.64898/2026.04.14.718476.

41. Sur, D., Agranyoni, O., Kirby, M., Cohen, N., Bagaev, A., Karandasheva, K., Shmerkin, E., Gorobets, D., Savita, B.K., Avneri, R., Divon, M.S., Lax, E., Michaelevski, I., and Pinhasov, A. (2023). Nurture outpaces nature: fostering with an attentive mother alters social dominance in a mouse model of stress sensitivity. Mol Psychiatry. 28(9):3816–3828. 10.1038/s41380-023-02273-y.

42. Walker, D.M., Cunningham, A.M., Gregory JK, Nestler EJ. (2019). Long-Term Behavioral Effects of Post-weaning Social Isolation in Males and Females. Front Behav Neurosci. 13:66. 10.3389/fnbeh.2019.00066.

43. Grigoryan GA, Pavlova IV, Zaichenko MI. (2022). Effects of Social Isolation on the Development of Anxiety and Depression-Like Behavior in Model Experiments in Animals. Neurosci Behav Physiol. 52(5):722–738. 10.1007/s11055-022-01297-1.

44. Yamamuro K, Bicks LK, Leventhal MB, Kato D, Im S, Flanigan ME, Garkun Y, Norman KJ, Caro K, Sadahiro M, Kullander K, Akbarian S, Russo SJ, Morishita H. (2020). A prefrontal-paraventricular thalamus circuit requires juvenile social experience to regulate adult sociability in mice. Nat Neurosci. 23(10):1240–1252. 10.1038/s41593-020-0695-6.

45. Einon DF, Morgan MJ. A critical period for social isolation in the rat. (1977) Dev Psychobiol. 10(2):123–32. 10.1002/dev.420100205.

46. Miczek KA, Barry H 3rd. (1975). What does the tube test measure? Behav Biol. 13(4):537–9. 10.1016/s0091-6773(75)91249-3

47. Reinwald JR, Ghanayem S, Wolf D, Scheller MF, Lebedeva J, Lebhardt P, Gölz O, Nelias C, Kelsch W. (2026) Individual differences drive social hierarchies in mouse societies eLife 15:RP109354. 10.7554/eLife.109354.1

48. Kappeler PM, Huchard E, Baniel A, Canteloup C, Charpentier MJE, Cheng L, Davidian E, Duboscq J, Fichtel C, Hemelrijk CK, Höner OP, Koren L, Micheletta J, Prox L, Saccà T, Seex L, Smit N, Surbeck M, van de Waal E and Girard-Buttoz C (2022) Sex and dominance: How to assess and interpret intersexual dominance relationships in mammalian societies. Front. Ecol. Evol. 10:918773. 10.3389/fevo.2022.918773.

49. Puentes-Escamilla, M.A., Schweinfurth, M.K. & Hemelrijk, C.K. (2024). When are females dominant over males in rats (*Rattus norvegicus*)?. Behav Ecol Sociobiol 78, 56. 10.1007/s00265-024-03465-4.

50. Yamaguchi, Y., Lee, Y.A., Kato, A., Jas, E., and Goto, Y. (2016). The Roles of Dopamine D2 Receptor in the Social Hierarchy of Rodents and Primates. Scientific reports, 7, 43348. 10.1038/srep43348.

51. Yamaguchi, Y., Lee, Y.A., Kato, A., and Goto, Y. (2017). The Roles of Dopamine D1 Receptor on the Social Hierarchy of Rodents and Nonhuman Primates. The international journal of neuropsychopharmacology, 20(4), 324–335. 10.1093/ijnp/pyw106.

52. Xing, B., Mack, N.R., Zhang, Y.X., McEachern, E.P., and Gao, W.J. (2022). Distinct Roles for Prefrontal Dopamine D1 and D2 Neurons in Social Hierarchy. Journal of Neuroscience, 42(2):313–324. 10.1523/JNEUROSCI.0741-21.2021.

53. Van der Kooij, M.A., Hollis, F., Lozano, L., Zalachoras, I., Abad, S., Zanoletti, O., Grosse, J., Guillot de Suduiraut, I., Canto, C., and Sandi, C. (2018). Diazepam actions in the VTA enhance social dominance and mitochondrial function in the nucleus accumbens by activation of dopamine D1 receptors. Molecular psychiatry, 23(3), 569–578. 10.1038/mp.2017.135.

54. Deng, X., Xu, W., Liu, Y., Jing, H., Zhong, J., Sun, K., Zhou, R., Xu, L., Wu, X., Zhang, B., Chen, W., Jiang, S., Chen, G., and Zhu, Y. (2025). Social rank modulates methamphetamine-seeking in dominant and subordinate male rodents via distinct dopaminergic pathways. Nature Neuroscience 28(6):1268–1279. 10.1038/s41593-025-01951-0.

55. Solié C, Nicolson A, Justo R, Layadi Y, Morin B, Batifol C, Reynolds LM, Le Borgne T, Fayad SL, Gulmez A, Rodriguez Quevedo Y, Allegret-Vautrot J, Centene Guglielmi G, de Chaumont F, Didienne S, Debray N, Hardelin JP, Girard B, Mourot A, Naudé J, Viollet C, Marti F, Delord B, Faure P. (2026). Dopaminergic mechanisms of dynamical social specialization. Nature. 10.1038/s41586-026-10301-4.

56. Grieb, Z.A., Lee, S., Stoehr, M.C., Horne, B.W., Norvelle, A., Shaughnessy, E.K., Albers, H.E., and Huhman, K.L. (2024). Sex-dependent regulation of social avoidance by oxytocin signaling in the ventral tegmental area. Behavioural Brain Research. 462:114881. 10.1016/j.bbr.2024.114881.

57. Bouarab, C., Wynalda, M., Thompson, B.V., Khurana, A., Cody, C.R., Kisner, A., and Polter, A.M. (2025). Sex-Specific Adaptations to VTA Circuits Following Subchronic Stress. European Journal of Neuroscience. 61(11):e70153. 10.1111/ejn.70153.

58. Wei A, Zhao A, Zheng C, Dong N, Cheng X, Duan X, Zhong S, Liu X, Jian J, Qin Y, Yang Y, Gu Y, Wang B, Gooya N, Huo J, Yao J, Li W, Huang K, Liu H, Mao F, Wang R, Shao M, Wang B, Zhang Y, Chen Y, Song Q, Huang R, Qu Q, Zhang C, Kang X, Xu H, Wang C. (2025). Sexually dimorphic dopaminergic circuits determine sex preference. Science. 387(6730):eadq7001. 10.1126/science.adq7001.

59. Shanley, M.R., Miura, Y., Guevara, C.A., Onoichenco, A., Kore, R., Ustundag, E., Darwish, R., Renzoni, L., Urbaez, A., Blicker, E., Seidenberg, A., Milner, T.A., and Friedman, A.K. (2023). Estrous Cycle Mediates Midbrain Neuron Excitability Altering Social Behavior upon Stress. Journal of Neuroscience. 43(5):736–748. 10.1523/JNEUROSCI.1504-22.2022.

60. Vandegrift, B.J., Hilderbrand, E.R., Satta, R., Tai, R., He, D., You, C., Chen, H., Xu, P., Coles, C., Brodie MS, and Lasek AW. (2020). Estrogen Receptor α Regulates Ethanol Excitation of Ventral Tegmental Area Neurons and Binge Drinking in Female Mice. Journal of Neuroscience 40(27):5196–5207. 10.1523/JNEUROSCI.2364-19.2020.

61. Rienecker, K.D.A., Chavasse, A.T., Moorwood, K., Ward, A., and Isles, A.R. (2020). Detailed analysis of paternal knockout Grb10 mice suggests effects on stability of social behavior, rather than social dominance. *Genes*, Brain and Behavior. 19(1):e12571. 10.1111/gbb.12571.

62. Seese, S., Tinsley, C.E., Wulffraat, G., Hixon, J.G., and Monfils, M.H. (2024). Conspecific interactions predict social transmission of fear in female rats. Scientific reports, 14(1):7804. 10.1038/s41598-024-58258-6.

63. Davidian E, Surbeck M, Lukas D, Kappeler PM, Huchard E. The eco-evolutionary landscape of power relationships between males and females. Trends Ecol Evol. 2022 Aug;37(8):706–718. doi: 10.1016/j.tree.2022.04.004. Epub 2022 May 18. PMID: 35597702.

64. Sapolsky, R.M. (2004). Social Status and Health in Humans and Other Animals. Annual Review of Anthropology 33: 393–418. 10.1146/annurev.anthro.33.070203.144000.

65. Sapolsky, R.M. (2005). The influence of social hierarchy on primate health. Science 308: 648–652. 10.1126/science.1106477.

66. Creel, S., Dantzer, B., Goymann. W., and Rubenstein DR. (2013). The ecology of stress: effects of the social environment. Functional Ecology 27: 66–80. 10.1111/j.1365-2435.2012.02029.x.

67. Cavigelli, S.A., and Caruso, M.J. (2015). Sex, social status and physiological stress in primates: the importance of social and glucocorticoid dynamics. Philos Trans R Soc Lond B Biol Sci 370(1669):20140103. 10.1098/rstb.2014.0103.

68. Sherman, G.D., Lee, J.J., Cuddy, A.J.C., Renshon, J., Oveis, C., Gross, J.J., and Lerner, J.S. (2012). Leadership is associated with lower levels of stress. Proceedings of the National Academy of Sciences 109: 17903–17907. 10.1073/pnas.1207042109.

69. von Rueden, C.R., Trumble, B.C., Emery Thompson, M., Stieglitz, J., Hooper, P.L., Blackwell, A.D., Kaplan, H.S., and Gurven, M. (2014). Political influence associates with cortisol and health among egalitarian forager-farmers. Evol Med Public Health 2014: 122–133. 10.1093/emph/eou021.

70. Williamson, C.M., Lee, W., Romeo, R.D., and Curley, J.P. (2017). Social context-dependent relationships between mouse dominance rank and plasma hormone levels. Physiology & behavior, 171, 110–119. 10.1016/j.physbeh.2016.12.038.

